# Chronic stress disrupts the network among stress granules, P-bodies, and motor proteins

**DOI:** 10.64898/2026.01.09.698697

**Authors:** Yuichiro Adachi, Masashi Masuda, Yutaka Taketani, Paul J. Anderson, Pavel V. Ivanov

## Abstract

Stress granules (SGs) are dynamic RNA granules that rapidly form in response to various stresses concurrent with mRNA translation shutdown, contributing to cellular adaptation and disease pathogenesis. While SG assembly and disassembly under acute stress have been extensively characterized, how SGs behave under chronic stress remains poorly understood. We previously reported that chronic stress preconditioning inhibits the earliest steps of SG assembly via translation-dependent mechanisms. In contrast, the regulation of SG maturation under chronic stress has not yet been investigated. Here, we show that chronic stress decreases SG size by disrupting the MYH9-dependent myosin crosstalk with core SG nucleator G3BP1. This defect leads to impaired SG and processing body (PB) docking as well, limiting the biogenesis of early SGs. Furthermore, chronic stress reduces expression of the SG nucleator UBAP2L, required for SG-PB docking, thus exacerbating these deficiences. In summary, chronic stress disrupts the myosin-SG-PB network and inhibits SG maturation in a translation-independent manner.

**Summary:** Chronic stress disrupts the MYH9 network with G3BP1, which inhibits stress granule (SG) assembly. It also decreases UBAP2L levels, which inhibits SG and processing body docking. Disrupted MYH9 network further inhibits this docking, thus blocking maturation of SGs.

## Introduction

To survive adverse environmental conditions, eukaryotic cells activate conserved stress-response pathways (Kim et al., 2008; Fulda et al., 2010). A central feature of the cellular stress response is protein synthesis shutdown (Pakos-Zebrucka et al., 2016), which reduces metabolic demands but paradoxically leads to the induction of the translation of some pro-survival, anti-apoptotic transcripts, which contribute to cytoprotection and damage repair. This general down-regulation of mRNA translation releases mRNAs and promotes the formation of transient cytoplasmic RNA biocondensates (Riggs et al. 2020). Stress granules (SGs) and processing bodies (PBs) are the best-characterized cytoplasmic RNA granules. SG assembly is initiated by polysome disassembly, which promotes release of untranslated mRNAs in the form of RNPs that interact with SG nucleators (e.g. G3BPs). PBs, in contrast, are constitutively present in the cytoplasm, but increase in number under cellular stress and often dock with SGs, potentially influencing SG size and maturation. (Buchan et al., 2008; Ripin et al., 2024; Riggs et al., 2024). SGs and PBs, while distinct cytoplasmic RNA granules, form a functional interface through extensive exchange of untranslated mRNAs and mRNPs under conditions of translational repression (Kedersha et al., 2005; Riggs et al., 2020). Proteins found in both SGs and PBs include tristetraprolin (TTP) and butyrate response factorlz1 (BRF1), which enhance physical interactions between SGs and PBs, as well as the caplzbinding initiation factor eIF4E, which is associated with capped mRNA (Kedersha et al., 2005). The shared mRNPs are accessible to components of both translational regulation and RNA decay pathways, thereby establishing a flexible network that modulates the fate of untranslated mRNAs (Decker and Parker, 2012).

Mechanistically, SG assembly is closely linked to the inhibition of mRNA translation initiation. Cellular stresses, such as an oxidative stress, an endoplasmic reticulum stress, or heat shock, activate the various kinases of the α-subunit of eukaryotic initiation factor 2 (eIF2α), leading to phosphorylation of eIF2α at Ser51 (Pakos-Zebrucka et al., 2016; Kedersha et al., 1999). Phosphorylated eIF2α inhibits the exchange of GDP for GTP on eIF2, thereby reducing the formation of the ternary complex required for delivery of initiator tRNA (tRNA_i_^Met^) to the 40S ribosomal subunit (Jackson et al., 2010). This globally suppresses translation initiation, causing polysomes to disassemble and liberating mRNAs that are no longer engaged in active translation. The released mRNAs, together with translation initiation factors, 40S ribosomal subunits, and RNA-binding proteins including G3BP1/2, undergo liquid–liquid phase separation (LLPS) to form SGs. Canonical SGs are membrane-less cytoplasmic RNA granules that contain 40S ribosome-contained mRNPs, mRNAs, RNA-binding proteins, and signaling factors. SG composition is stress-specific and plays a decisive role in directing cell fate toward survival or death (Ivanov et al., 2019; Aulas et al., 2017). For instance, while sodium arsenite (SA), most frequently used drug to mimic oxidative stress, promotes the canonical SG assembly and leads to cell survival, nitric oxide induces the assembly of eIF3b-lacking non-canonical SGs and promotes cell death (Aulas et al., 2018). SGs contribute to the excessive aggregation of disease-related proteins with neurodegenerative diseases such as amyotrophic lateral sclerosis (ALS), Alzheimer’s disease (AD), frontotemporal dementia (FTD), etc (Wolozin and Ivanov, 2019). PBs are cytoplasmic RNA granules enriched in translationally repressed mRNAs and mRNA decay factors and are organized by core scaffolding proteins such as Hedls/EDC4, EDC3, and the RNA helicase DDX6, which coordinate mRNA storage and turnover (Kedersha et al., 2005; Buchan and Parker, 2009; Riggs et al., 2020). Through these components, PBs function as hubs that couple translational repression to mRNA storage or decay under cellular stress (Andrei et al., 2005; Decker and Parker, 2012; Hubstenberger et al., 2017). Dysregulation of PB components and PB dynamics has been implicated in human disease, including cancer and viral infections (Anderson et al., 2015; Kanakamani et al., 2021).

During SG biogenesis, various physical interactions between different SG components occurs, some of which promote gathering of SG “seeds” into small-sized SGs, and their further fusion into larger microscopically-visible SGs. Actin and microtubules play complementary roles in the positioning of SGs in the cytosol (Böddeker et al., 2022), and the dynamics of SGs are also proposed to be influenced by cytoskeleton-associated motor proteins (Panas et al., 2016). The increase of SG size to microscopically visible SGs is due to fusion of smaller SGs, in turn promoted by the transportation along microtubules by their associated motor proteins (Nadezhdina et al., 2010; Chernov et al., 2009). Importantly, pharmacological and genetic experiments revealed that disruption of microtubule arrays interferes with SG formation without robust effect on protein synthesis (Ivanov et al., 2003; Loschi et al., 2009). This suggests that network of microtubules-motor proteins contributes to SG assembly. On the other side, the actin filament network may also contribute to SG biogenesis and dynamics. Specific motor proteins such as non-muscle myosin II isoform A (NMMHC- II A) and myosin Va are reported to influence assembly and size of SGs under acute stress (Hu et al., 2023; Zhou et al., 2024). Yet, molecular details and physiological conditions leading to the decoupling between cytoskeleton, motor protein and SGs have not been reported. During SG biogenesis, the ultimate size and maturation of SGs are determined not only by cytoskeleton-mediated transport and fusion of smaller SG seeds, but also by opposing activities at the SG-PB interface. On the PB side, the RNA helicase DDX6 acts to restrict SG-PB docking, thereby limiting the expansion of SGs (Ripin et al., 2024), whereas on the SG side, UBAP2L promotes condensation and accelerates docking with PBs, facilitating SG growth (Riggs et al., 2024). However, the mechanistic link between cytoskeleton-mediated SG transport and PB interactions, and the contribution of specific modulators to SG size control, remains unclear.

Not only acute stress but also chronic stress contributes to the progression of various diseases, including neurodegenerative disorders and cancer (Kim et al., 2008; Fulda et al., 2010; Wolozin and Ivanov, 2019). While SGs are rapidly assembled in response to acute stress as part of the integrated stress response (ISR) with the phosphorylation of eIF2α to protect cells from cytotoxic insults, studies investigating the effects of chronic stress on SG dynamics remain limited. Pre-conditioning is generally defined as the phenomenon where exposure to low levels of stress do not cause severe cell damage but could protect against subsequent exposure to more severe stress. These pre-conditioned chronic stress leads to eIF2α phosphorylation and activates pro-survival pathways with global translation shutdown to conserve cellular energy and production of cytoprotective molecules (like ATF4) (Lu et al., 2004; Peng et al., 2012). Nevertheless, we recently reported that the preconditioned cell fails to form SGs in response to the later acute SA stress, ultimately leading to excessive cell death without the canonical SG-mediated cytoprotection. Chronic stress slows down mRNA. translation elongation, which consequently impedes the disassembly of polysomes and the condensation of G3BPs and mRNPs. It should be noted however that excessively high doses of SA incubation can overcome such block and disassemble polysomes completely, promoting SG formation even under chronic stress, yet with delayed onset (Adachi et al., 2025). The detailed nature of these SGs and the molecular mechanisms of SG assembly under chronic stress still lack many details.

Here, we describe how chronic stress impairs the maturation of SGs by disrupting their fusion. The fusion is regulated by cytoskeleton-associated motor proteins, and interactions with PBs. We show that chronic stress specifically decreases the binding between G3BP1 and MYH9, a central protein of non-muscle myosin IIA, which leads to block in the transition of small sized SGs into microscopically visible SGs, and larger SGs. We also report that the MYH9 network plays a role not only in SG assembly but also in SG-PB interactions. Chronic stress induces the reduction of UBAP2L levels, which leads to less efficient SG-PB docking and inhibits SG maturation, potentially promoting excessive disruption of the MYH9 network.

## Results

### Chronic stress restricts SG assembly and maturation

To understand the characters of SG formation in chronically stressed cells, we preincubated human osteosarcoma U2OS cells with a low dose of sodium arsenite (SA) (10 µM) for 24 h, which did not induce SG assembly, prior to treating with various dose of SA (50, 100, 200, 300, 400, and 500 µM) (Fig. 1). We have reported chronic stress (10 µM SA for 24 h) reduces the SG positive cells by additional 50 and 100 µM SA stress (Adachi et al., 2025), but only 200 µM of SA-induced SG formation was also inhibited by chronic stress and higher doses of SA (300, 400, 500 µM) was not affected regarding SG-positive cells (Fig. 1 A–C). Moreover, chronic stress decreases SG-positive cells and the average number of SGs only under lower doses of additional SA (SG positive cells: 50, 100, 200 µM, average SG number: 100, 200, 300 µM), in contrast, chronic stress pre-conditioned cells demonstrate significantly smaller in size SGs at any tested doses (Fig. 1 C–E). Importantly, chronic stress by other types of stresses than oxidative stress, such as ER stress (Tg), mitochondrial stress (CCCP), and starvation stress (HBSS), also decreased the SG positive cells, average number of SGs, and the size of SGs (Sup Fig. 1). We also tested the effects of excessive stress on pre-conditioned cells in a time-dependent manner (0-240 min) (Sup Fig. 2). SG appearance at earlier time points (30 – 45 min of SA 500 µM incubation) was evidently delayed by chronic stress (Sup Fig. 2 B), same as the previous report (Adachi et al., 2025). The number and size of SGs in pre-conditioned cells were smaller at most of the time points (Sup Fig. 2 C–D). In addition, wash-out experiments from 500 µM SA incubation did not show any differences in the formation and size of SGs (Sup Fig. 3). Taken together, we conclude that chronic stress specifically inhibits the assembly and maturation of SGs but not their disassembly and degradation.

**Figure 1.**
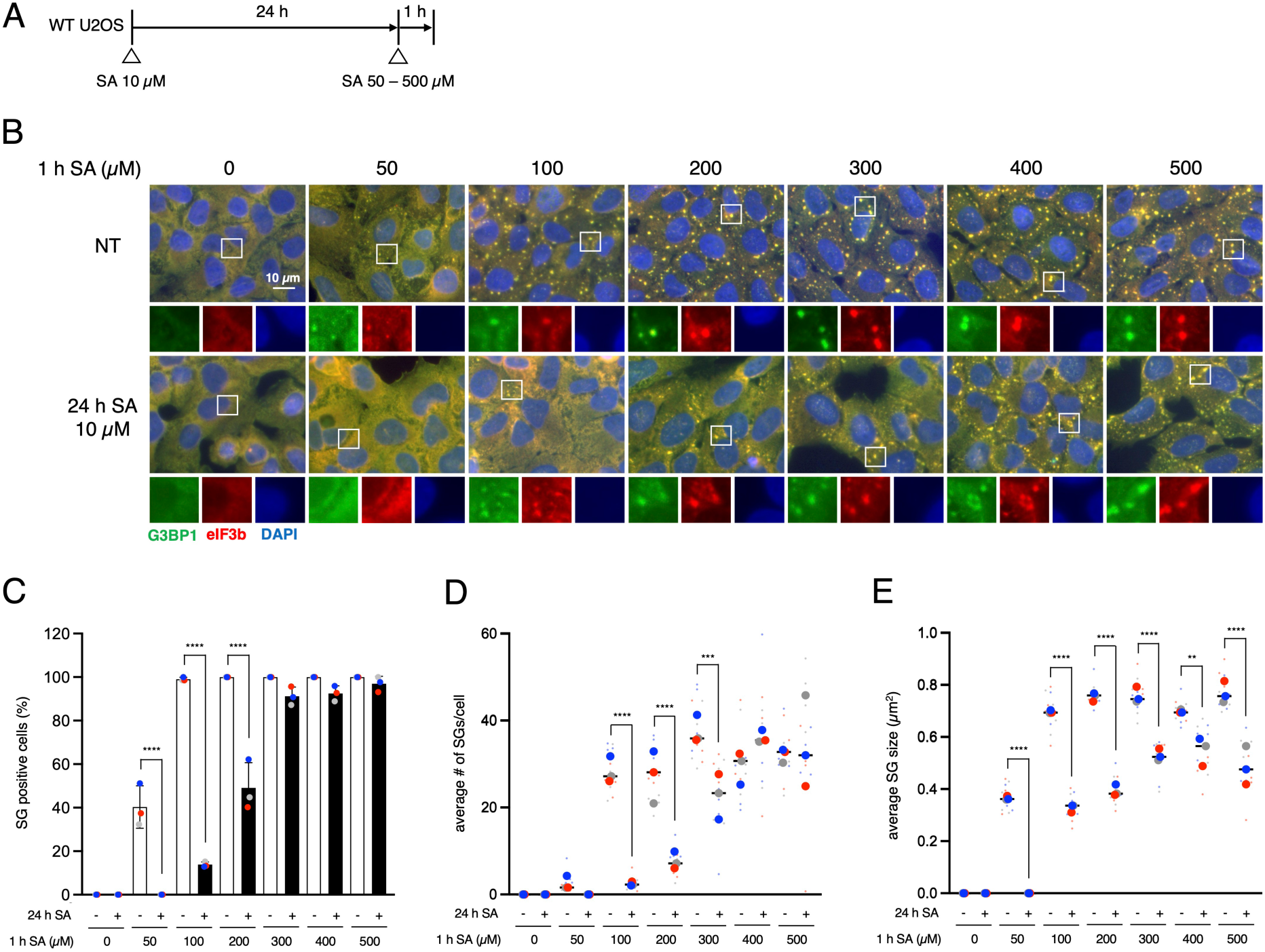
Chronic stress interferes with SG formation. (A) Schematic illustration of the experimental timeline. (B–E) U2OS cells were subjected to treatment with 50, 100, 200, 300, 400 or 500 µM SA for 1 h after preincubation with 10 µM SA for 24 h. Unstressed cells (NT) were used as a control. (B) Representative images of U2OS cells stained with G3BP1 (green), eIF3B (red), and DAPI (blue). Scale bar = 10 µm. (C) Cells were examined for the presence of the core SG markers G3BP1, and eIF3B. (D) Cells were examined for the average number of G3BP1 per cell. (E) Cells were examined for the average size of G3BP1. (C–E) Four images were taken for each of the three biological replicates (independent experiments). Large circle dots represent the mean results from the images (n = 4 technical replicates) of one biological replicate, the same colors of dots are from the same independent experiment. Small circle dots in different shades of the same color are from the same independent experiment. P values were assessed using a one-way ANOVA (p** < 0.01, p*** < 0.001, p**** < 0.0001). Results are mean ± S.E.M. (n = 3).

**Figure 2.**
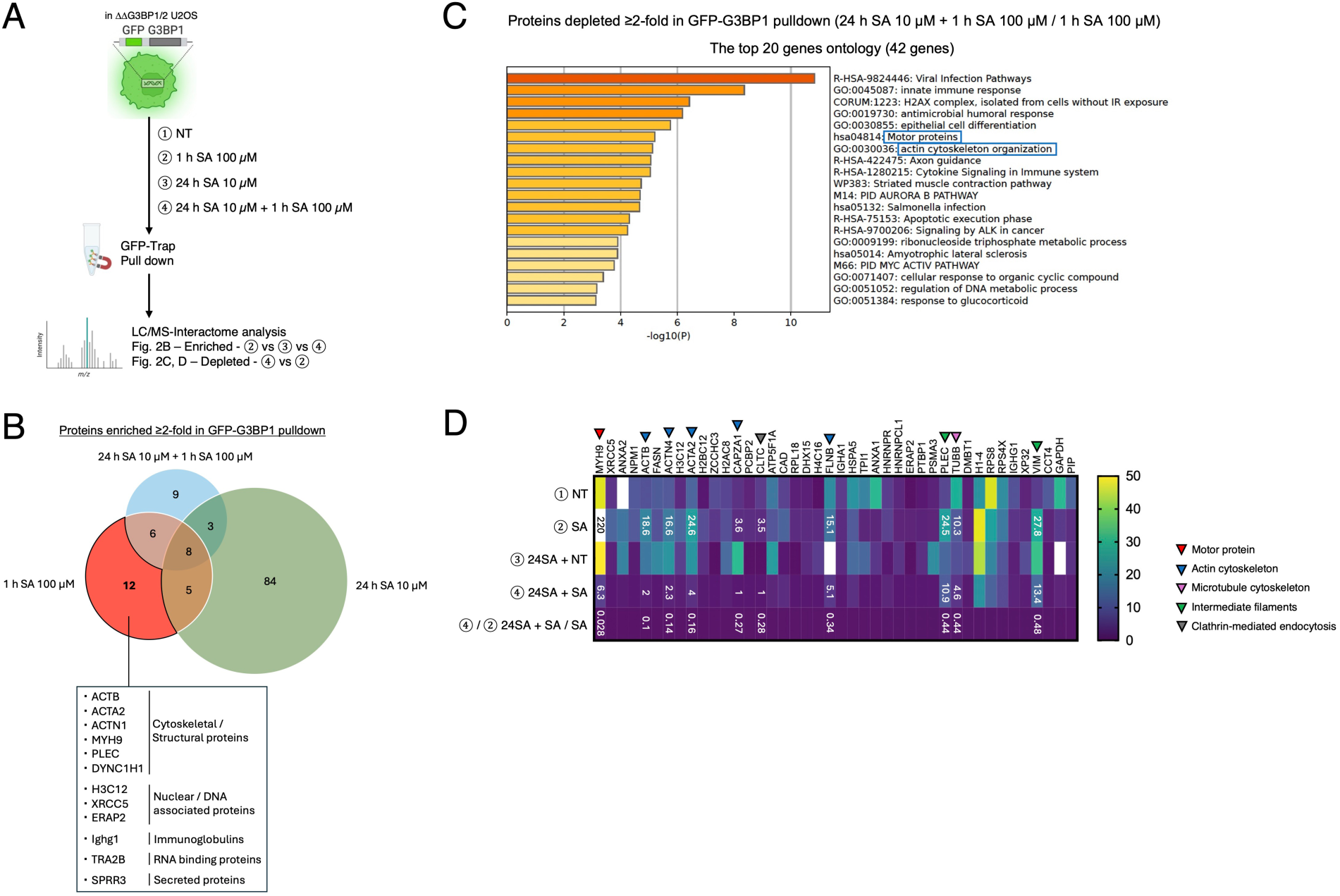
Chronic stress induces unique G3BP1-Binding Interactome patterns. (A–D) GFP-G3BP1-ΔΔG3BP1/2-U2OS and GFP-U2OS cells were subjected to treatment with 100 µM SA for 1 h after pre-incubation with 10 µM SA. Lysates were applied to GFP-trap pull-down and Mass spectrometry analysis. (A) Schematic illustration of the experimental timeline. (B) Venn diagram of the >2-fold enriched proteins (vs. NT) in GFP-G3BP1 GFP-trap pull-down. (C–D) The comparison between ④ 24 h SA 10 µM + 1 h SA 100 µM and ② 1 h SA 100 µM. (C) Top 20 depleted pathways (down-regulated) by G3BP1-Binding Interactome. (D) The list of the most down-regulated 42 candidates.

**Figure 3.**
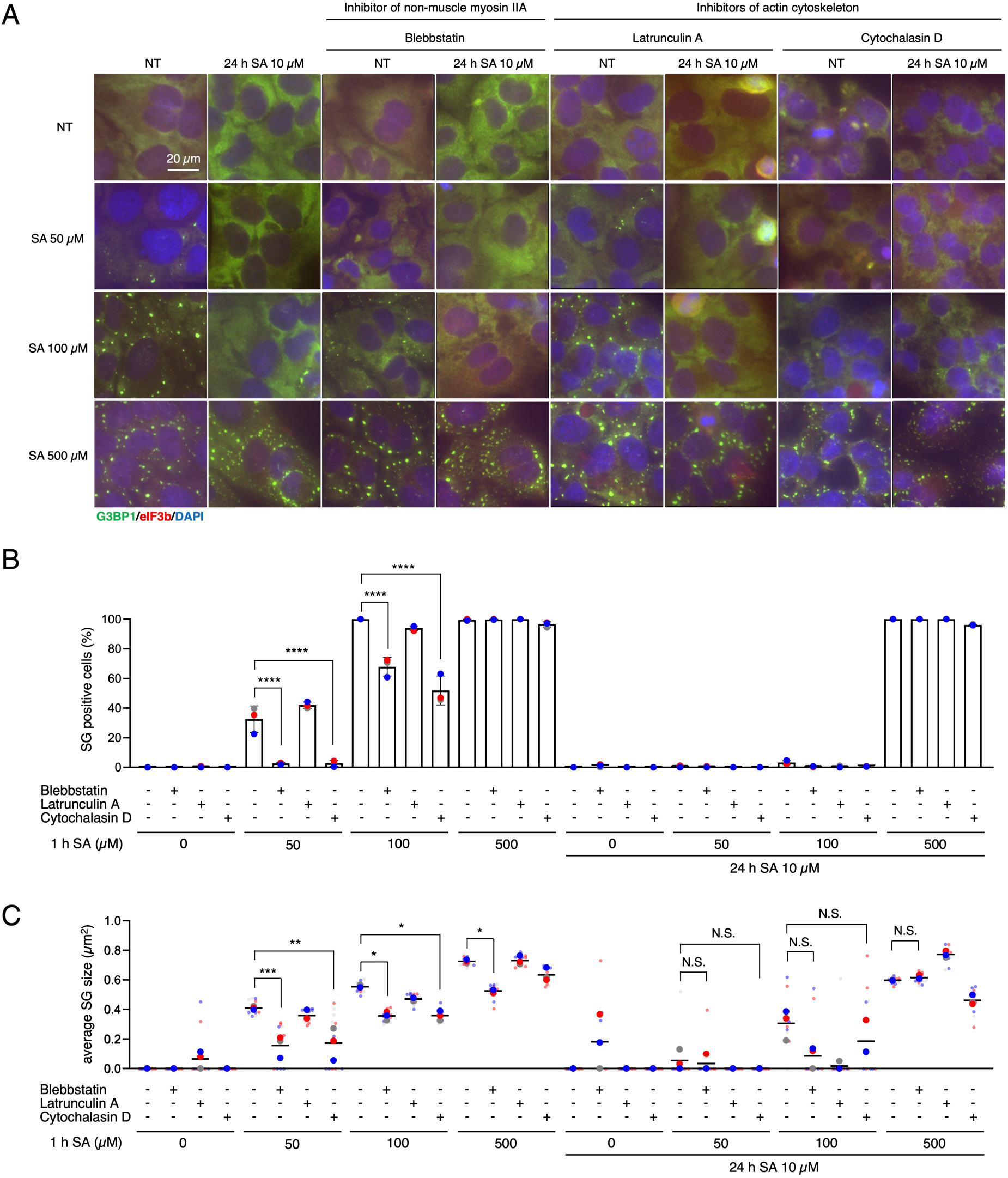
Chronic stress promotes formation of smaller SGs through the disruption of non-muscle myosin IIA. (A–C) U2OS cells were subjected to treatment with 50, 100, or 500 µM SA for 1 h after preincubation with 10 µM SA and/or 100 µM blebbstatin, 0.2 µM latrunculin A, and 0.5 µM cytochalasin D for 24 h. Unstressed cells (NT) were used as a control. (A) Representative images of U2OS cells stained with G3BP1 (green), eIF3B (red), and DAPI (blue). Scale bar = 20 µm. (B) Cells were examined for the presence of the core SG markers G3BP1, and eIF3B. (C) Cells were examined for the average size of G3BP1. (B–C) Four images were taken for each of the three biological replicates (independent experiments). Large circle dots represent the mean results from the images (n = 4 technical replicates) of one biological replicate, the same colors of dots are from the same independent experiment. Small circle dots in different shades of the same color are from the same independent experiment. P values were assessed using a one-way ANOVA (p* < 0.05, p** < 0.01, p*** < 0.001, p**** < 0.0001). Results are mean ± S.E.M. (n = 3).

### Chronic stress disrupts the network between the SG proteome and cytoskeletal proteins

To identify the mechanisms of the interruption of SG assembly and maturation, we probed the proteome-interactome of G3BP1 through mass spectrometry analysis in response to different SA regime stresses (Fig. 2). From the Venn diagram that shows the enriched proteins in GFP-G3BP1 pulldown, cytoskeletal/structural proteins were shown as a top pathway by acute SA stimulation, and those where not shown in chronic stress groups (Fig. 2 B). Gene ontology analysis comparison between cells after acute 1 h SA 100 µM stress and chronic 24 h SA 10 µM stress followed by acute stress revealed that chronically stressed cells demonstrated overall less association with motor proteins and actin cytoskeleton organizations (Fig. 2 C). Motor proteins, actin-cytoskeleton, and microtubules are proposed to control the cytosolic position of SGs, affecting their dynamics and regulating their size (Böddeker et al., 2022; Panas et al., 2016; Nadezhdina et al., 2010; Chernov et al., 2009). MYH9, the heavy chain of non-muscle myosin of class II isoform A (NM IIA), was the top candidate in our analysis to reveal the most reduced interaction with G3BP1 by acute 1 h SA with chronic 24 h SA stress. Multiple actin cytoskeleton-related proteins were also shown in that list, as well as one of the microtubule cytoskeleton-related proteins. There are unique unknown proteins as candidates of SG regulators, intermediate filaments, and clathrin-mediated endocytosis (Fig. 2 D). This interactome analysis suggests the possibility that chronic stress results in immature SGs because of the reduced interaction between G3BPs and these motor proteins, cytoskeleton-related factors, and other unique candidates.

### Disruption of non-muscle myosin IIA, but not microtubule-related cytoskeleton, by chronic stress results in smaller SGs

Since G3BP1 interactome analysis under chronic stress showed reduced binding to several specific protein groups, including motor proteins and cytoskeletal components, we decided to use various cytoskeleton-related inhibitors, with or without chronic stress preincubation, to determine which protein(s) are primarily associated with formation of smaller SGs under chronic stress. We firstly preincubated cells with 10 µM SA and/or blebbstatin, a chemical inhibitor of NM IIA, latrunculin A, and cytochalasin D, inhibitors of the actin cytoskeleton, prior to treating with 50, 100, and 500 µM SA for 1 h (Fig. 3). Blebbstatin is known to reduce the number and the size of SGs induced by 43 ℃ heat shock, but nothing has been reported in a SA dose-dependent manner (Hu et al., 2023). Blebbstatin itself inhibited both the SG-positive cells and the SG size with 50 and 100 µM of SA. Interestingly, blebbstatin did not change the percentage of SG-positive cells but decreased the size of SG even with 500 µM SA (Fig. 3 B–C), which indicates NM IIA affects SG maturation rather then the formation of visible SG. Importantly, chronic SA stress decreases SG size, but co-incubation with blebbstatin did not change SG size (Fig. 3C). Blebbstatin incubation also did not affect translation efficiency and translation was completely shut off under 500 µM SA stress (Sup Fig. 4). This suggests chronic stress inhibits the maturation of SGs via the disruption of NM IIA network, not polysome disassembly. Latrunculin A sequesters G-actin monomers to prevent actin polymerization, whereas cytochalasin D binds to the barbed ends of F-actin, blocking filament elongation and promoting filament disruption; both ultimately lead to disruption of the actin cytoskeleton. Latrunculin A reduces the number of SGs (Ivanov et al., 2003), but there are no reports about latrunculin A and cytochalasin D on the effects of SG dynamics. Surprisingly, latrunculin A did not affect any SG characters (Fig. 3 B–C). Cytochalasin D decreased the percent of SG-positive cells and SG size with 50 and 100 µM of SA, like blebbstatin, however, it did not affect the 500 µM SA-induced SG characteristics with and without chronic stress (Fig. 3 B–C).

**Figure 4.**
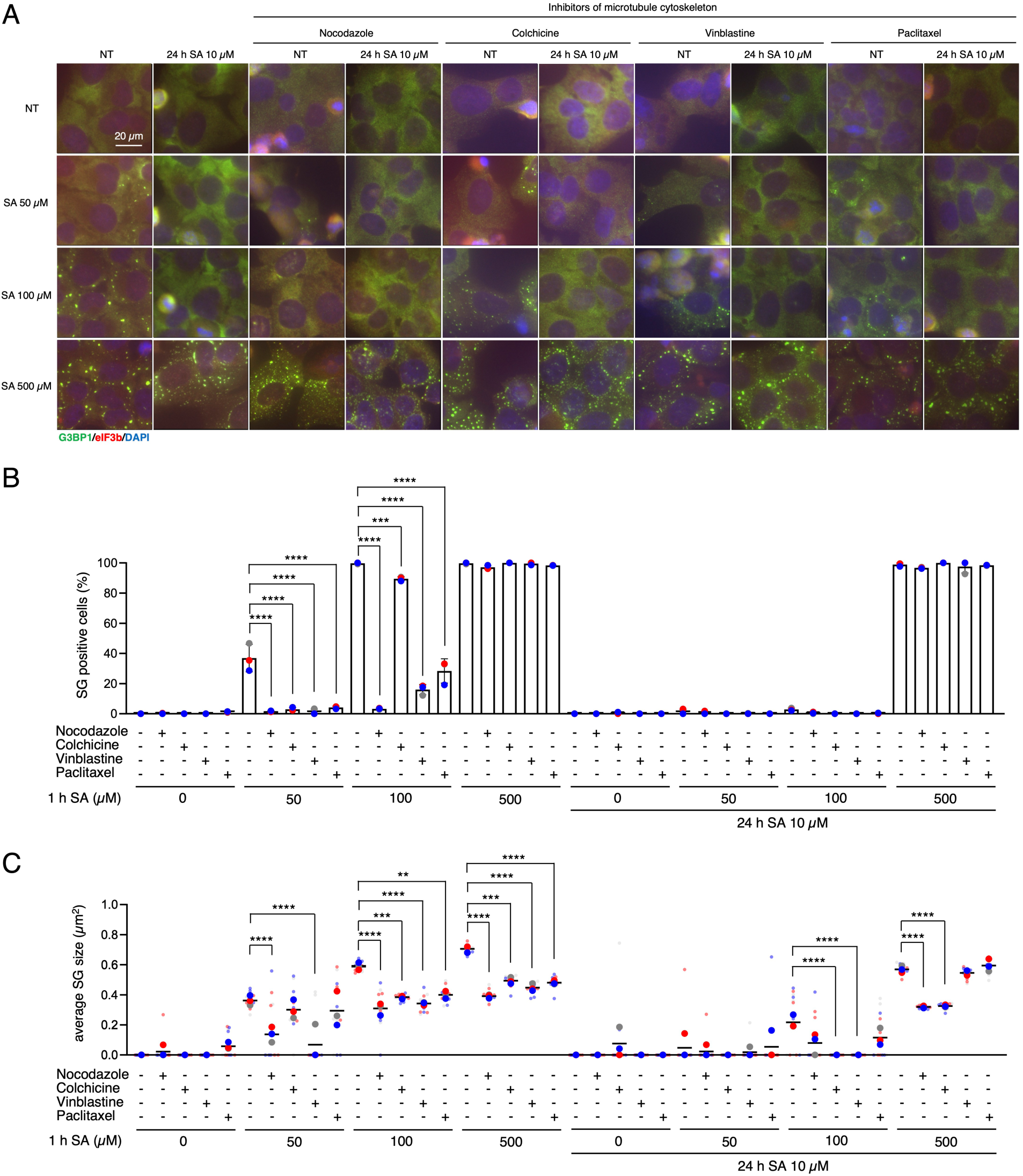
Microtubule cytoskeleton regulates formation and size of SGs but is not associated with chronic stress changes. (A–C) U2OS cells were subjected to treatment with 50, 100, or 500 µM SA for 1 h after preincubation with 10 µM SA and/or 0.1 µM nocodazole, 5 nM colchicine, 50 nM vinblastine, and 50 nM paclitaxel for 24 h. Unstressed cells (NT) were used as a control.= (A) Representative images of U2OS cells stained with G3BP1 (green), eIF3B (red), and DAPI (blue). Scale bar = 20 µm. (B) Cells were examined for the presence of the core SG markers G3BP1, and eIF3B. (C) Cells were examined for the average size of G3BP1. (B–C) Four images were taken for each of the three biological replicates (independent experiments). Large circle dots represent the mean results from the images (n = 4 technical replicates) of one biological replicate, the same colors of dots are from the same independent experiment. Small circle dots in different shades of the same color are from the same independent experiment. P values were assessed using a one-way ANOVA (p** < 0.01, p*** < 0.001, p**** < 0.0001). Results are mean ± S.E.M. (n = 3).

Next, we tested the effects of nocodazole, colchicine, vinblastine, and paclitaxel, inhibitors of microtubule cytoskeleton, on the formation and the size of SGs (Fig. 4). Nocodazole and colchicine inhibit microtubule polymerization by binding to tubulin, whereas vinblastine induces microtubule depolymerization and paclitaxel stabilizes microtubules by preventing their disassembly; all of these agents disrupt microtubule dynamics and thereby inhibit microtubule function. It has been reported that all three inhibitors—nocodazole, vinblastine, and paclitaxel—except colchicine, reduce the number of SG-positive cells (Ivanov et al., 2003). In our data, all of these reagents decreased the percentage of SG-positive cells by 50 and 100 µM, yet 500 µM SA-induced SGs were not affected. Nevertheless, all inhibitors decreased the size of SGs with any dose of SA. Nocodazole and colchicine, both inhibitors of microtubule polymerization, decreased the size of SGs even under chronic stress, whereas two other inhibitors did not affect it (Fig. 4 B–C). Given the inconsistent results, we propose that SG maturation failure under chronic stress is largely independent of microtubule disruption.

We’ve also examined the unique hits as decreased binding with G3BP1 under chronic stress, using withaferin A, an inhibitor of intermediate filaments, and pitstop 2, an inhibitor of Clathrin-mediated endocytosis (Sup Fig. 5). Previous proteomic analysis showed that withaferin A increases the G3BP1 expression and promotes SG formation (Kumar et al., 2021), but there are no reports on relationship of pitstop 2 with SG formation. Unlike previous reports, long-time and low doses of withaferin A did not promote SG formation. Moreover, it significantly decreased the percentage of SG-positive cells with 100 µM SA. However, withaferin A increased the number of SG-positive and size of SGs in chronically stressed cells. Pitstop 2 also increased the SG-positive cells and SG size (Sup Fig. 5 B–C).

**Figure 5.**
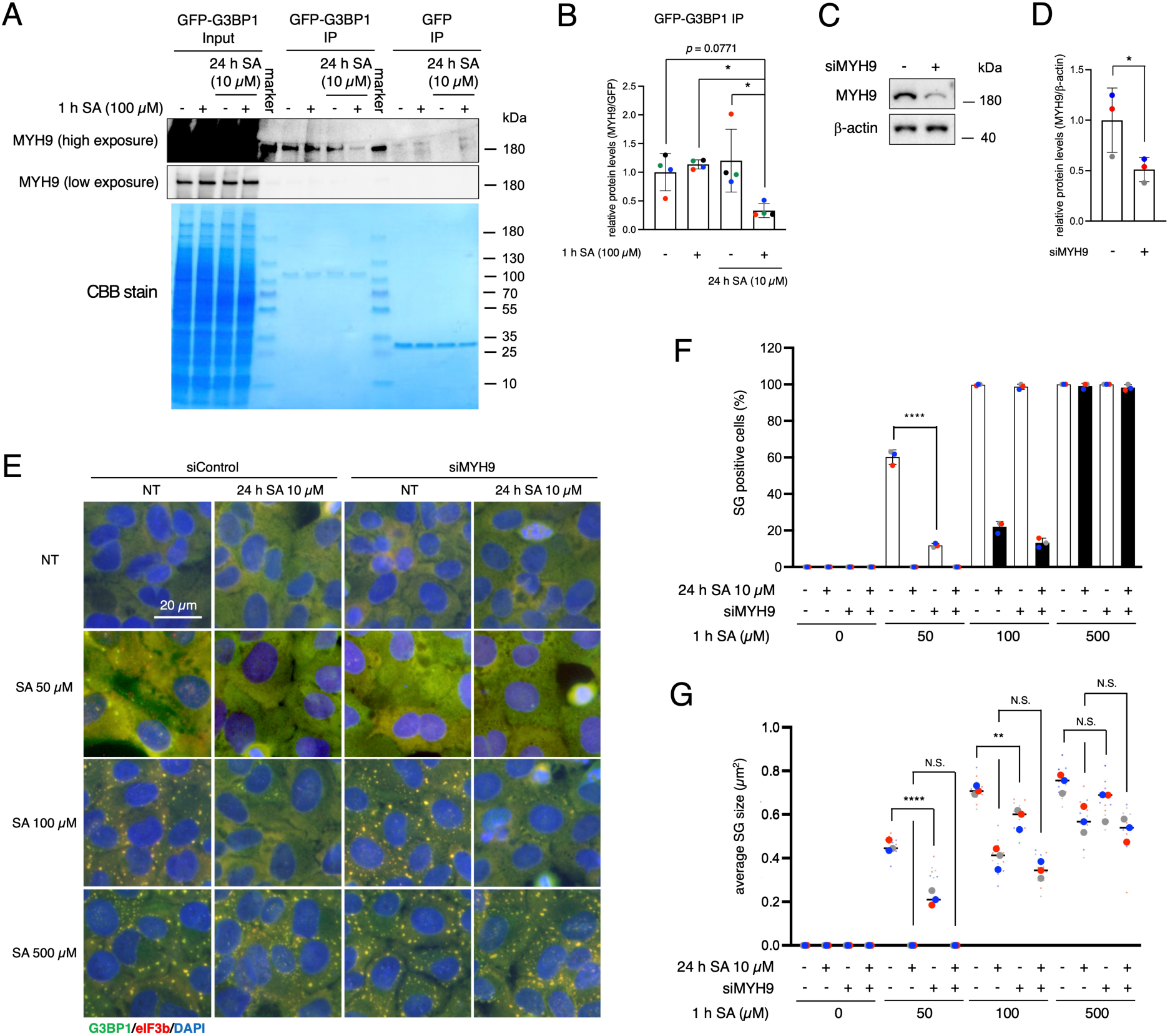
Chronic stress disrupts binding between G3BP1 and MYH9. (A) Cell lysates were subjected to western blotting using antibodies for MYH9, and CBB stained. (B) Proteins were quantified from blots using densitometry and normalized to GFP. Four blots were taken for each of the four biological replicates (independent experiments). The same colors of the circle dots in the graph are from the same independent experiment. P values were assessed using a one-way ANOVA (p* < 0.05). Results are mean ± S.E.M. (n = 4). (C–G) U2OS cells were treated with siRNA against MYH9 or non-targeting siRNA. (E–G) siRNA-treated U2OS cells were subjected to treatment with 50, 100, or 500 µM SA for 1 h after preincubation with 10 µM SA. Unstressed cells (NT) were used as a control. (C) Cell lysates were subjected to western blotting using antibodies for MYH9 and β-actin. (D) Proteins were quantified from blots using densitometry and normalized to β-actin. Three blots were taken for each of the four biological replicates (independent experiments). The same colors of the circle dots in the graph are from the same independent experiment. P values were assessed using an unpaired two-tailed t-test (p* < 0.05). Results are mean ± S.E.M. (n = 3). (E) Representative images of blebbstatin-treated U2OS cells stained with G3BP1 (green), eIF3B (red), and DAPI (blue). Scale bar = 20 µm. (F) Cells were examined for the presence of the core SG markers G3BP1, and eIF3B. (G) Cells were examined for the average size of G3BP1. (F–G) Four images were taken for each of the three biological replicates (independent experiments). Large circle dots represent the mean results from the images (n = 4 technical replicates) of one biological replicate, the same colors of dots are from the same independent experiment. Small circle dots in different shades of the same color are from the same independent experiment. P values were assessed using a one-way ANOVA (p** < 0.01, p**** < 0.0001). Results are mean ± S.E.M. (n = 3).

Our proteomic analysis suggested that G3BP1 interactome was significantly decreased under chronic stress followed by acute SA insult (Fig. 2). Consequently, we also probed the MYH9-bound levels with G3BP1 by immunoprecipitation assay. Western blotting confirmed that MYH9-G3BP1 binding decreased specifically when cells underwent acute stress after chronic stress (10 µM SA for 24 h +1 h of 100 µM SA) (Fig. 5 A–B). To determine whether a decrease in the interactions between G3BP1 and MYH9 contributes to this phenomenon, we used siRNA to deplete MYH9 and confirmed the effects of the MYH9 network on the formation and size of SGs (Fig. 5 C–G). siMYH9 significantly decreased MYH9 expression levels (Fig. 5 C–D). MYH9 knockdown itself decreased both the percentage of SG-positive cells and the size of SGs with 50 µM SA, but only SG size was decreased under 100 µM SA. This is similar to the results with blebbstatin under 500 µM SA incubation, suggesting that NM IIA network has important roles in SG assembly and maturation. Notably, this siMYH9 effects were less efficient than chronic stress. 500 µM SA-induced SGs were not affected by the knockdown of MYH9 (Fig. 5 E–G). Considering MYH9 levels were approximately half compared with the basal condition by siRNA (Fig. 5 D), and whole NM IIA inhibitor blebbstatin inhibited SG maturation but not as potently as by chronic stress, more severe MYH9 knockdown or knockout may affect SG regulation even with 500 µM SA. Taken together, we concluded NM IIA including MYH9 may play a role in the biogenesis of smaller SGs and maturation of SGs.

### Chronic stress and the disruption of myosin IIA inhibit of SG-PB docking

PBs frequently interact with SGs, and this interaction has been implicated in the regulation of SG growth and maturation (Kedersha et al., 2005; Riggs et al., 2024). However, the contribution of SG-PB docking to chronic stress-induced immature SG inductions has not been clarified. We also preincubated cells in the same way with 10 µM SA for 24 h as a chronic stress, prior to treating with 50, 100, and 500 µM SA for 1 h, and checked the localization of Hedls/EDC4, a typical PB marker, with G3BP1 (Fig. 6). The size of SGs was decreased by chronic stress (Fig. 6 B), same as previously. Interestingly, the percentage of docking PBs to SGs was also decreased by chronic stress (Fig. 6 C). Since chronic stress did not change the numbers and the size of PBs under most of the SA doses (Fig. 6 D–E), chronic stress may direcly block the docking of SGs and PBs and, thus, reduce the SG size. Taken together with the possibility that NM IIA plays a role in the maturation of SGs (Fig. 3 and 5), we preincubated cells with 10 µM SA and/or blebbstatin, before treating with 50, 100, and 500 µM SA for 1 h (Fig. 6 F–I). The size of SGs was reduced by blebbstatin itself and that effect was further enhanced by chronic stress (Fig. 6 G), same as previously (Fig. 3 B). In addition, the percentage of docking PBs was also limited by blebbstatin (Fig. 6H). Although chronic stress accelerated the reduction of docking PBs by blebbstatin under 500 µM SA (Fig. 6H), the average number of PBs was also decreased (Fig. 6G), which further limits PBs for docking with SGs. Considering that chronic stress-induced reduction in SG size was unaffected by blebbstatin, the changes in SG–PB docking under chronic stress may also occur through disruption of the MYH9 network.

**Figure 6.**
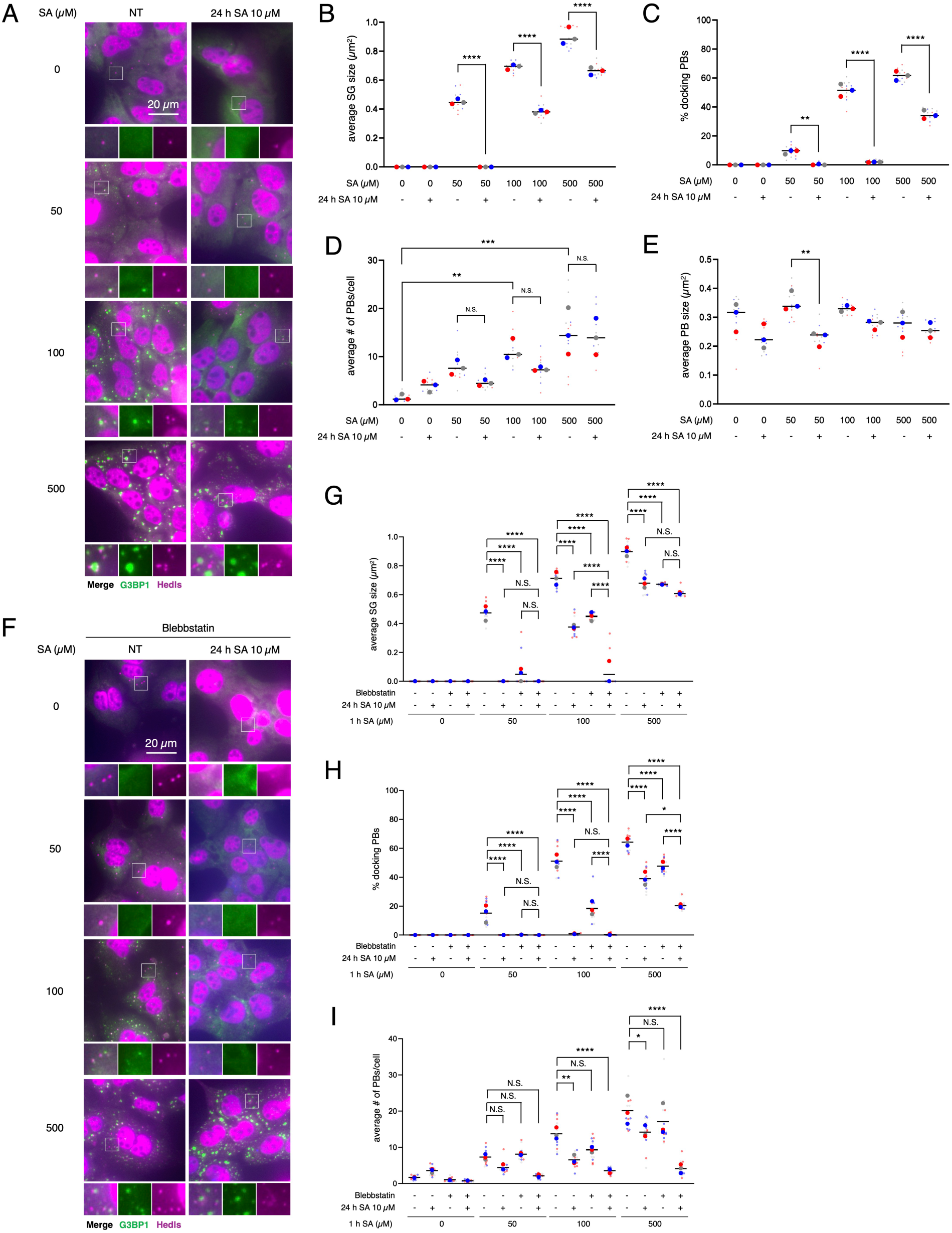
Chronic stress negatively regulates docking between P-bodies and SGs. (A–D) U2OS cells were subjected to treatment with 50, 100, or 500 µM SA for 1 h after preincubation with 10 µM SA. Unstressed cells (NT) were used as a control. (A) Representative images of U2OS cells stained with G3BP1 (green), Hedls (purple), and DAPI (blue). Scale bar = 20 µm. (B) Cells were examined for the average size of G3BP1. (C) Cells were examined for the % of PBs (Hedls) docking to SGs (G3BP1). (D) Cells were examined for the average number of Hedls per cell. (E) Cells were examined for the average size of Hedls. (F–I) U2OS cells were subjected to treatment with 50, 100, or 500 µM SA for 1 h after preincubation with 10 µM SA and/or 100 µM blebbstatin. (F) Representative images of U2OS cells incubated with blebbstatin stained with G3BP1 (green), Hedls (purple), and DAPI (blue). Scale bar = 20 µm. (G) Cells were examined for the average size of G3BP1. (H) Cells were examined for the % of PBs (Hedls) docking to SGs (G3BP1). (I) Cells were examined for the average number of Hedls per cell. (B–E, G–I) Four images were taken for each of the three biological replicates (independent experiments). Large circle dots represent the mean results from the images (n = 4 technical replicates) of one biological replicate, the same colors of dots are from the same independent experiment. Small circle dots in different shades of the same color are from the same independent experiment. P values were assessed using a one-way ANOVA (p* < 0.05, p** < 0.01, p*** < 0.001, p**** < 0.0001). Results are mean ± S.E.M. (n = 3).

### Chronic stress disrupts the network of SGs, PBs, and MYH9 through UBAP2L reduction

We further examined the expression levels of PB/SG-related proteins in the chronically stressed cells. Interestingly, the expression levels of Ubiquitin Associated Protein 2 Like (UBAP2L), a core SG factor that cooperates with G3BPs to promote SG formation and interactions with PBs, were significantly decreased by SA chronic stress (Fig. 7 A–B). This reduction of UBAP2L was not observed at early time points of SA incubation (1, 4 h) and occurred in a time-dependent manner (Sup Fig. 6 A). We also confirmed this UBAP2L reduction not only in chronic SA incubation but also in other stresses (Sup Fig. 6 B), which reduced the size of SGs (Sup Fig 1). We then preincubated both WT and ΔUBAP2L U2OS cells in the same way with 10 µM SA for 24 h as a chronic stress, prior to treating with 50, 100, and 500 µM SA for 1 h (Fig. 7 C–G). Same as reported previously (Riggs et al., 2024), ΔUBAP2L cells showed smaller SGs and fewer SG-PB dockings with any dose of acute SA incubation (Fig. 7 D–E). Although the number and size of SGs were not changed by chronic stress in ΔUBAP2L cells (Fig. 7 F–G), the SG size and docking PBs under 500 µM SA acute stress were decreased by chronic stress (Fig. 7 D–E), indicating that while UBAP2L is essential for SG–PB docking and the regulation of SG size, severe depletion of UBAP2L could exacerbate the blockade of SG maturation, even during chronic stress. To examine another consequence of UBAP2L reduction, we assessed MYH9 expression, and notably, MYH9 protein levels were reduced in ΔUBAP2L U2OS (Fig. 8 A). Surprisingly, the depletion of not only UBAP2L but also the G3BPs decreased the MYH9 levels (Fig. 8 B). MYH9 levels were not changed by each of the only single knockout of G3BPs suggesting their redundancy (Fig. 8 C) and as reported before (Kedersha et al., 2016). We also confirmed that effects of G3BPs’ depletion on MYH9 expression are replicated in other cell line, namely breast cancer MDA-MB-468 cells (Fig. 8 D). These indicate that SG core proteins stabilize MYH9 network, and/or vice versa, and chronic stress may disrupt this network by both by limiting their interactions and reduction in expression of MYH9 proteins. Altogether, our data suggest that chronic stress preconditioning interferes with the MYH9 network, which then inhibits SG assembly and SG-PB docking, thus limiting SG maturation. Chronic stress also reduces the UBAP2L expression and down-regulates docking of PBs, resulting in decrease of SG size (Fig. 9 A).

**Figure 7.**
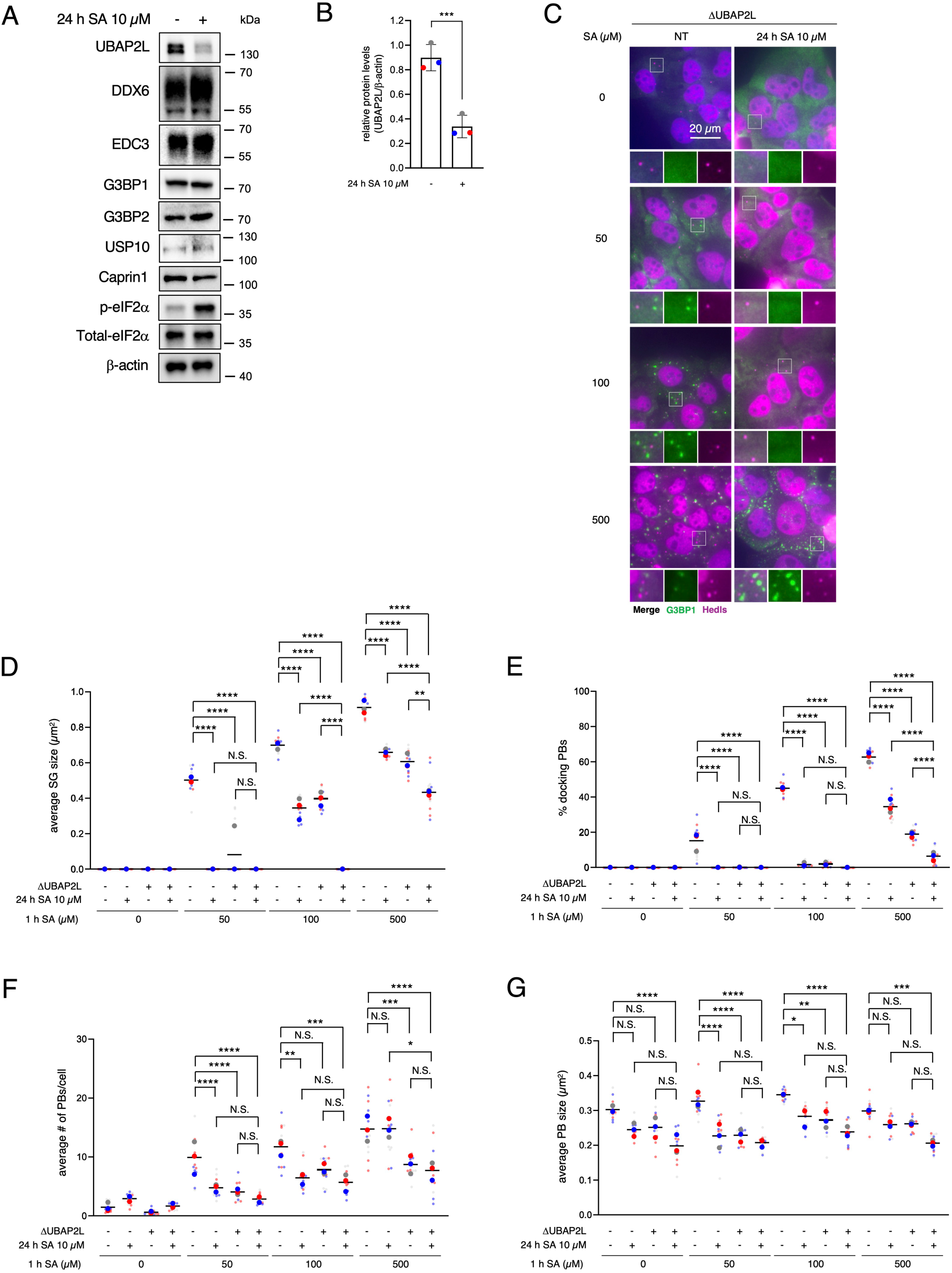
Chronic stress decreases UBAP2L expression levels and inhibits the docking between P-bodies and SGs. (A–B) U2OS cells were subjected to treatment with 10 µM SA for 24 h. Unstressed cells (NT) were used as a control. (A) Cell lysates were subjected to western blotting using antibodies for UBAP2L, DDX6, EDC3, G3BP1, G3BP2, USP10, Caprin1, p-eIF2α, total eIF2α, and β-actin. (B) Proteins were quantified from blots using densitometry and normalized to β-actin. Three blots were taken for each of the four biological replicates (independent experiments). The same colors of the circle dots in the graph are from the same independent experiment. P values were assessed using an unpaired two-tailed t-test (p* < 0.05). Results are mean ± S.E.M. (n = 3). (C–G) WT and ΔUBAP2L U2OS cells were subjected to treatment with 50, 100, or 500 µM SA for 1 h after preincubation with 10 µM SA. Unstressed cells (NT) were used as a control. (C) Representative images of ΔUBAP2L U2OS cells stained with G3BP1 (green), Hedls (purple), and DAPI (blue). Scale bar = 20 µm. (D) Cells were examined for the average size of G3BP1. (E) Cells were examined for the % of PBs (Hedls) docking to SGs (G3BP1). (F) Cells were examined for the average number of Hedls per cell. (G) Cells were examined for the average size of Hedls.

**Figure 8.**
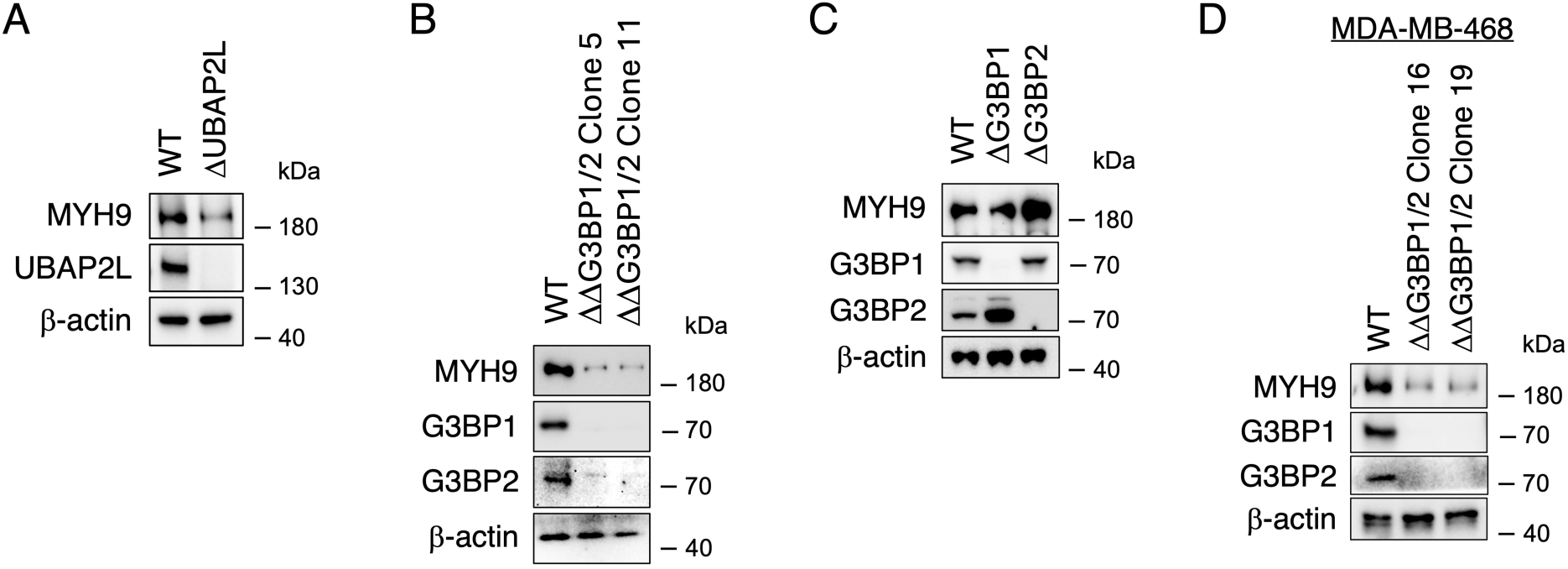
Depletion of G3BPs and UBAP2L destabilize MYH9 protein. (A) WT and ΔUBAP2L U2OS cells were lysed and subjected to western blotting using antibodies for MYH9, UBAP2L, and β-actin. (B–D) Cell lysates were subjected to western blotting using antibodies for MYH9, G3BP1, G3BP2, and β-actin. (B) WT, ΔΔG3BP1/2 (Clone 5 and 11) U2OS cells (C) WT, ΔG3BP1, and ΔG3BP2 U2OS cells. (D) WT, ΔΔG3BP1/2 (Clone 16 and 19) MDA-MB-468 cells.

**Figure 9.**
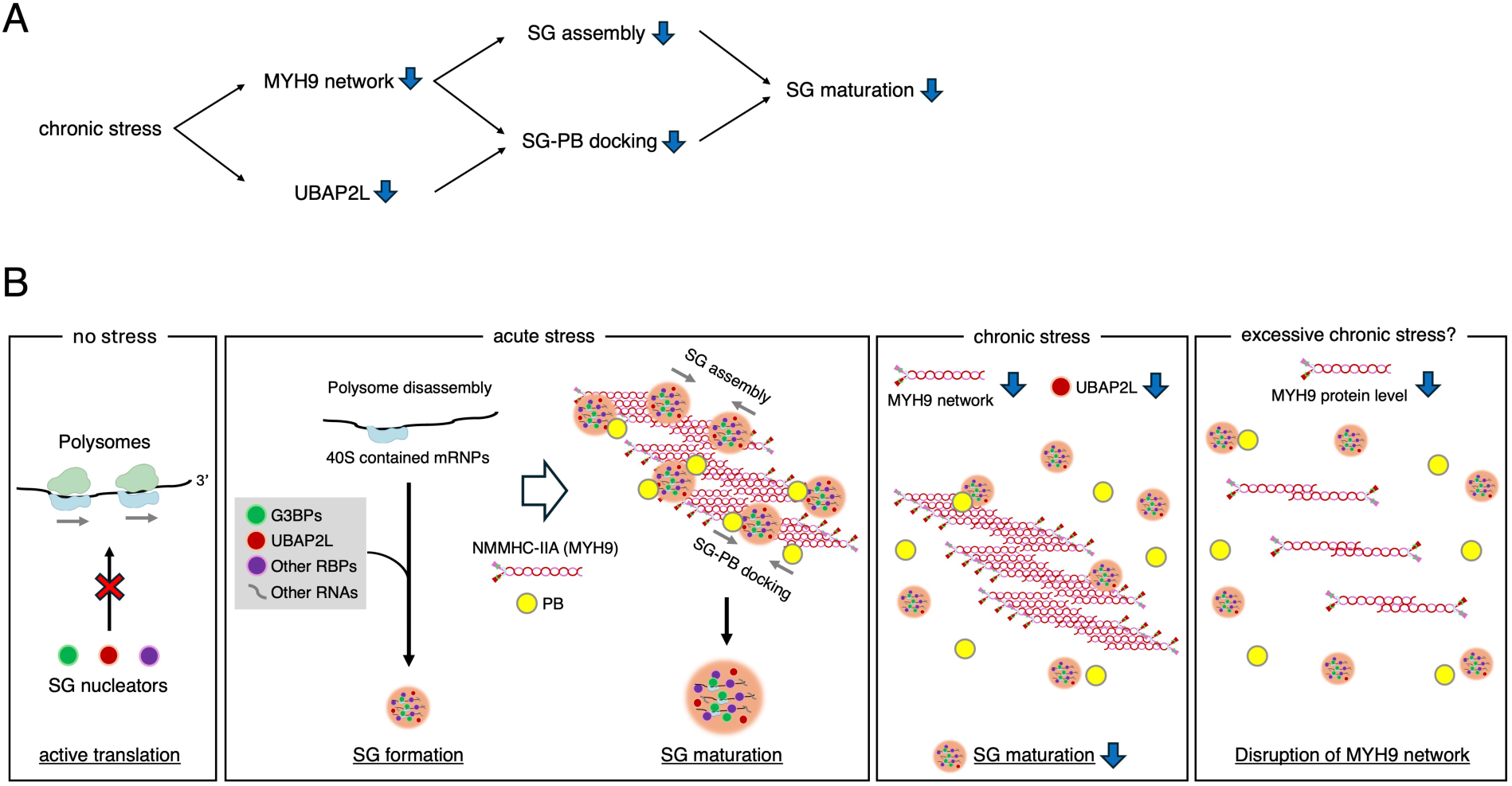
Schematic illustration of the disrupted SG-P-body-MYH9 network in chronic stress pre-incubated cells. (A) Schematic summary of the effects of chronic stress on SG maturation: Chronic stress interferes with the MYH9 interaction with G3BP1, thus inhibiting SG assembly, and decreases UBAP2L protein expression levels, which induces less efficient SG-PB docking. Disruption of the MYH9 network also inhibits SG-PB docking. Decrease in both SG assembly and SG-PB docking lead to immature SGs. (B) Proposed schematic model of SG maturation failure under chronic stress: Active polysomes block mRNP condensation with SG nucleators under no stress conditions. Acute stress disassembles polysomes and induces SGs with nucleation of SG-related RBPs, including G3BPs and UBAP2L. Once SG seeds are formed, SGs are further assembled through the interaction between G3BPs and MYH9. SG and PB docking is also promoted by the non-muscle myosin II isoform A (NMMHC-IIA). Both SG assembly and SG-PB docking lead to SG maturation and promote formation of larger SGs. Chronic stress decreases the interaction of G3BPs and MYH9 and down-regulates protein levels of UBAP2L, which is the core factor of SG-PB docking. Since NMMHC-IIA also promotes SG-PB docking, chronic stress inhibits SG maturation through both the weak MYH9 network (less SG assembly via G3BPs and less SG-PB docking) and fewer SG-PB dockings with decreased UBAP2L. Taken together, the depletion of G3BPs or UBAP2L decreases MYH9 protein levels, while chronic stress itself reduces UBAP2L expression; therefore, excessive chronic stress may induce a profound disruption of the MYH9 network.

## Discussion

SG assembly and disassembly in response to acute stress have been broadly characterized, reflecting their ability to rapidly sense and respond to cellular stress (Wheeler et al., 2016). In contrast, SG dynamics under chronic stress have been less investigated, even though SGs play critical roles in the pathophysiology of neurodegenerative diseases, cancer, inflammatory disorders, and other conditions associated with prolonged stress (Wolozin and Ivanov, 2019; Riggs et al., 2020; Li and Wang, 2023; Ratti et al., 2020). Others and we have reported the effects of chronic stress on SG formation. However, there are limited reports of long-term exposure to stress or drug-induced SGs: chronic glucose starvation induces pro-death SGs, 24 h of cisplatin incubation induces SGs (Reineke et al., 2010; Martin et al., 2023), and the effects of inhibition of SG formation: chronic proteasomal inhibition impairs SG formation, mitochondrial unfolded protein response-induced GADD34 inhibits SG formation (Shelkovnikova et al., 2017; Lopez-Nieto et al., 2024). These researchers suggested that the impaired SG formation is due to insufficient eIF2α phosphorylation. However, we have reported that even under chronic stress, there can be enough eIF2α phosphorylation (Fay et al., 2021), and SG assembly is inhibited due to ribosome stalling associated with chronic stress-induced eEF2 phosphorylation (Adachi et al., 2025). Our current study indicates other mechanistic ways to interfere with SG formation, not related to translation regulation. NM II is an actin-associated motor, and actin filaments contribute to the movement of RNA granules related to SGs and PBs in plant cells (Hamada et al., 2012), while in mammals movement of PBs and SGs was shown to be microtubule-dependent (Bartoli et al., 2011; Loschi et al., 2009). Our proteomic data and results based on the use of NM II complex inhibitor blebbstatin suggest that chronic stress also affects physical interaction between G3BP and the subunit of NM II, MYH9. As NM II was shown to be a constituent of early SGs (Hu et al., 2023), we suggest that disruption of G3BP-MYH9 interaction prevents coalescence of smaller earlier SGs into larger ones through both SG seeds assembly and SG-PB contacting.

SG composition varies in a stress and time-specific manner, exerting diverse effects on intracellular homeostasis (Aulas et al., 2017). In addition to influencing cell fate decisions such as cell survival and protection (Ivanov et al., 2019; Aulas et al., 2017), SGs have been shown to contribute to fatty acid β-oxidation, lysosomal damage responses, and stress-responsive signaling pathways including MAPK and caspases, etc (Bussi et al., 2023; Amen and Kaganovich, 2021; Arimoto et al., 2008; Fujikawa et al., 2023; Duran et al., 2024). It is suggested that SG maturation has several stages, and the different SG sizes may have different dynamics (Protter and Parker, 2016). If the size of SGs affects the patterns of SG components and regulates SG function, SGs in chronically stressed cells may have a unique function (loss –or gain-of-function). Interestingly, we also found out that withaferin A, an inhibitor of intermediate filaments, and pitstop 2, an inhibitor of Clathrin-mediated transport, could rescue the diminished size of SGs by chronic stress (Sup Fig. 5 C). These unique findings may represent an important resource for future efforts to develop disease therapies targeting SGs. Previous studies showed that various inhibitors of myosin, cytoskeleton, and microtubule interfere with SG maturation (Ivanov et al., 2003; Loschi et al., 2009; Hu et al., 2023). It should also be noted that higher concentrations of vinca alkaloids (high micromolar concentrations) such as vinblastine or vincristine paradoxically promote SG formation and promote protein synthesis shutdown (Szaflarski et al., 2016). Our study reports new data suggesting that low-dose various stress-inducible reagents disrupt the myosin network.

Interestingly, UBAP2L expression level was decreased by chronic stress, but not by acute stress (Fig. 7 A–B and Sup Fig. 6). There are no reports to show the steady or inducible decrease of UBAP2L expression except for specific genetic diseases. Since UBAP2L is a key SG nucleator, and depletion of UBAP2L decreases SG formation (Yang et al., 2020; Cirillo et al., 2020), it is possible that continuous chronic stress reduces the efficiency of SG formation and SG-PB docking, simultaneously. This may be connected to etiology of some UBAP2L-associated genetic diseases such as UBAP2L-deficiency syndrome, a rare autosomal dominant neurodevelopmental disorder caused by loss-of-function mutations in UBAP2L, characterized by impaired language development, intellectual disability, behavioral abnormalities, and dysmorphic facial features (Jia et al., 2022; Yang et al., 2024). Since SGs play critical roles in maintaining neuronal homeostasis (Wolozin and Ivanov, 2019), and both genetic and stress-induced impairment of UBAP2L converge on defective SG dynamics, SG-UBAP2L axis may share molecular mechanisms contributing to neurological phenotypes.

MYH9 expression, meanwhile, is reported to decrease not only as a result of genetic alterations but also under conditions of chronic stress in the kidney: glomerular MYH9 expression is repressed by HIV-1 infection, and it’s also suppressed in diabetic patients/animal/podocytes treated with angiotensin 2 (Hays et al., 2012; Kang et al., 2019). While much of the SG research has focused on neurodegenerative diseases and cancer, a remarkable finding is that acute oxidative stress induces SG formation in the kidney (yet chronic stress has not been examined), as reported in models of metabolic and ischemic renal stress (Wang et al., 2019; Zhu et al., 2025). MYH9 is also known to be regulated by protein levels through degradation by the ubiquitin-proteasome pathway, and the direct interaction of deubiquitinase YOD1 stabilizes MYH9 protein (Sun et al., 2024). SGs are RNA biocondensates formed through liquid-liquid phase separation (LLPS) and proteins that drives LLPS typically contain intrinsically disordered regions (IDRs) (Molliex et al., 2015). Both G3BP1 and UBAP2L have IDRs (Hofmann et al., 2021), but interestingly, approximately half of the amino acids in MYH9 are also predicted to contain IDRs (Sup Fig. 7). As depletion of UBAP2L or G3BPs reduced MYH9 expression (Fig. 8), MYH9 may interact with these SG core proteins and be stabilized and perform its function correctly via IDRs. ΔUBAP2L cells show low expression levels of MYH9 (Fig. 8 A). In contrast, 24 h SA chronic oxidative stress did not decrease the MYH9 levels, even though it lowers UBAP2L levels (Fig. 5 A and Fig. 7 A–B). Taken together, excessive chronic stress may indirectly reduce MYH9 expression through UBAP2L downregulation, potentially disrupting cytoskeletal integrity and SG dynamics.

In conclusion, as summarized in Fig. 9 B, we propose the model of chronic stress-induced SG maturation failure through both defects in SG assembly and SG-PB docking caused by a weak MYH9 network. Chronic stress ultimately leads to MYH9 network disruption via UBAP2L reduction. SGs dynamics and their regulation have been studied extensively and discussed the potential contribution to especially neurodegenerative disease (Wolozin and Ivanov, 2019; Advani and Ivanov, 2020), however, our research uncovered a novel interaction between RNA granules and motor proteins through various chronic stresses. This implies the possibility of more broad contribution of SG biology to various diseases.

## Materials and methods

### Cell culture and drug treatment

Human osteosarcoma U2OS cells and Triple-negative breast cancer MDA-MB-468 cells (Table S1) were maintained at 37 °C in a 5.0% CO2 in DMEM (Corning) containing 20 mM HEPES (Gibco), 10% FBS (Sigma), 100 U/ml penicillin, and 100 µg/ml streptomycin. Genetically modified cells were previously established through CRISPR-Cas9 technology (Kedersha et al., 2016; Sanders et al., 2020; Yang et al., 2024 *preprint*) (Table S1). For any experiments, cells were grown to ~70% confluency and then treated as indicated in figure legends: sodium arsenite (SA, Sigma), thapsigargin (Tg, Invitrogen), carbonyl cyanide 3-chlorophenylhydrazone (CCCP, Sigma), HBSS +calcium +magnesium (HBSS, Gibco), blebbstatin (Cayman), latrunculin A (Cayman), cytochalasin D (Cayman), nocodazole (Cayman), colchicine (Cayman), vinblastine (Cayman), paclitaxel (Cayman), withaferin A (Cayman), and pitstop 2 (Cayman).

### siRNA knockdowns

Approximately 100,000 cells were seeded per well in 6-well plates overnight. Cells were transfected with 50 nM siRNA (Table S2) with 2.5 µl Lipofectamine 2000 (Invitrogen). 24 h after transfection, cell culture medium was replaced with fresh medium. At 48 h, cells were transfected with a second dose of siRNA, as at t = 0. At ~72 h, cells were counted and reseeded in 12-well plates (~200,000 cells/well) and 24-well plates (~100,000 cells/well) for immunostaining and western blot analysis, respectively. At ~96 h cells in 24-well plates were stress-treated (if desired) and harvested for immunofluorescence or western blotting.

### Immunofluorescence

Cells were plated into a 24-well plate seed with coverslips. The following day, cells were treated as indicated in figure legends. Then the cells were fixed with 4% paraformaldehyde for 15 min, permeabilized with −20 °C methanol for 5 min, and blocked for 1 h with 5% normal horse serum (NHS; ThermoFisher) diluted in PBS. Primary antibodies shown in Table S3 were diluted in blocking solution and incubated for 1 h at room temperature or overnight at 4 °C. Next, cells were washed three times and then incubated with secondary antibodies (Jackson Laboratories) and Hoechst 33258 (Sigma-Aldrich) for 1 h at room temperature and washed. Coverslips were mounted on glass slides with Vinol and imaged.

### Microscopy

Wide-field fluorescence microscopy was performed using an Eclipse E800 microscope (Nikon, Minato, Tokyo, Japan) equipped with epifluorescence optics and a digital camera (Spot Pursuit USB) for Fig. 1 and Sup Fig. 1. Images were merged using Adobe Photoshop. Eclipse Ni-E microscope (Nikon, Minato, Tokyo, Japan) and NIS-Elements imaging software (Nikon, Minato, Tokyo, Japan) were used for all images except those above. Image acquisition was done with a 40× objective (PlanApo; Nikon, Minato, Tokyo, Japan). Analysis was performed using FIJI. Samples were split into channels, converted to grayscale, and each stack was combined into one image comprised of the intensity of each slice summed together. This allowed us to capture the entirety of the signal from the foci of interest. The threshold tool was used to create ROIs for the foci of interest and the cytoplasm. G3BP1 was used to create the SG ROI. Hedls was used to create PB ROI. This analysis was performed on four images per condition across three biological replicates, from which the mean and standard error of the mean were calculated.

### SG, PB, and docking quantification

SG and PB counts, as well as their docking, were quantified in U2OS cells using a FIJI macro originally described by Riggs C et al., 2024. Briefly, nuclei were excluded based on Hoechst staining, SGs were detected by top-hat filtering and MaxEntropy thresholding, and PBs were detected by Gaussian subtraction followed by MaxEntropy thresholding. Masks of SGs and PBs were then used to identify overlapping areas representing docking events.

### GFP Trap pull-down assay and Pathway/Venn diagram analysis

GFP-G3BP1-ΔΔG3BP1/2-U2OS and GFP- ΔΔG3BP1/2-U2OS cells were washed twice with ice-cold 1X PBS, and lysed with 1X IP buffer (Cell Signaling, #9803). ChromoTek GFP-Trap® Magnetic Agarose bead (Proteintech, # gtma) was washed twice with ice-cold cell 1X IP buffer (Cell Signaling, #9803) by Magnetic stand. Each cell supernant lysate was incubated with the pre-washed GFP- Trap® Magnetic bead at 4°C for 1 hour with shaking on a rotator. Unbound proteins were removed by washing 3 times with the IP Buffer, and proteins bound to GFP-Trap® Magnetic bead were eluted with SDS loading buffer. The analysis of binding partners of G3BP1 was performed by Taplin Mass Spectrometry Facility in Harvard Medical School. Gene ontology (GO) enrichment and Kyoto Encyclopedia of Genes and Genomes (KEGG) pathway enrichment analyses were performed with Metascape (https://metascape.org/), an online resource for integrated gene list annotation and analysis (Zhou et al., 2019). Quantitative Venn diagrams were created using DeepVenn (https://www.deepvenn.com/).

### Western blotting

Following drug treatment, cells were washed with phosphate-buffered saline (PBS). Protein samples were heated to 95°C for 10 min in Laemmli sample buffer in the presence of 100 mM dithiothreitol (DTT) and subjected to SDS–PAGE. Samples were loaded on a 4–20% Tris-Glycine gel (BioRad) and transferred to nitrocellulose membrane. Membranes were blocked with Tris-buffered saline with 0.1% Tween-20 (TBS-T) with 5% milk for 1 h at room temperature. Antibodies were diluted in 5% normal horse serum in PBS. Primary antibodies were incubated overnight at 4°C and secondary antibodies for 1 h at room temperature. Antibody information is listed in Table S3. Antibody detection was performed using SuperSignal West Pico Chemiluminescent Substrate (Thermo Scientific). Gels were stained with GelCode™ Blue Safe Protein Stain (CBB stain, Thermo Fisher Scientific) for 1 hour at room temperature and subsequently destained with deionized water until protein bands were clearly visible.

### RiboPuromycylation assay

Ribopuromycylation assay was described in Panas et al., 2015. In brief, cells were unstressed or stressed as indicated. 5 min before harvest, puromycin and emetine were added to a final concentration of 9 and 91 µM, respectively, and the incubation continued for 5 min. Cells were then lysed and subjected to Western blotting using an anti-puromycin antibody.

### IDR prediction

Intrinsically disordered regions (IDRs) of human MYH9 were analyzed using the PRDOS prediction tool (https://prdos.hgc.jp/cgi-bin/top.cgi) according to the server’s default parameters. The results are presented in Supplementary Figure 7.

### Statistics, reproducibility and data analysis

Means were compared to each other using One-way ANOVA or an unpaired two-tailed t test. Results are mean values ± standard deviation (SD) of at least three independent experiments, or results show one representative experiment of a minimum of three biological replicates. Statistical analyses were performed on all available data. Statistical tests were performed using GraphPad Prism software. SGs and PBs were visualized by IF with indicated markers. For quantifications, four fields were taken from selected samples, with three replicated independent experiments. Cells were considered SG-positive if they have at least three cytoplasmic foci.

## Supporting information

Mas spec data

## Data availability

The data are available from the corresponding author upon reasonable request. The mass spectrometry data are shown in Table S4. Raw mass spectrometry data of proteins for Fig. 2. Sheet 1: raw data. Sheet 2: proteins enriched >2 by 1 h SA 100 *µ*M. Sheet 3: proteins enriched >2 by 24 h SA 10 *µ*M. Sheet 4: proteins enriched >2 by 24 h SA 10 *µ*M + 1 h SA 100 *µ*M. (Sheet 2–4 is for Fig. 2 A) Sheet 5: proteins depleted >2 by 24 h SA 10 *µ*M + 1 h SA 100 *µ*M compared with 1 h SA 100 *µ*M. (for Fig. 2 C) Sheet 6: values of counted peptides for Fig. 2 D.

## Acknowledgments

We thank members of Ivanov and Anderson labs for the valuable discussion. Especially, we sincerely thank Claire Riggs (Present address: St. Olaf College, Northfield, MN, USA) for her help in understanding the code related to SG–PB docking. The figures are created with BioRender.com. This work was supported by funds from National Institutes of Health grant R35 GM126901 (P.J.A.), National Institutes of Health grant R01 GM126150 and R01 GM146997 (P.I.), Japan Society for the Promotion of Science (JSPS) Grants-in-Aid for Scientific Research (PS KAKENHI) 22KJ2354 (Y.A.), 20K21761 and 21H03359 (M.M.), 19H04053 and 23H03329 (Y.T.). Y.A. was supported by the JSPS Overseas Challenge Program for Young Researchers.

