## Supplementary Information for "Chronic stress disrupts the network among stress granules, P-bodies, and motor proteins"

**Table S1.** Cell lines used in this study and their sources.

| Cell line | Source |
| --- | --- |
| U2OS-WT | ATCC |
| U2OS-G3BP1/G3BP2 KO double (DDG3BP1/2) | (Kedersha et al., 2016) |
| U2OS-GFP-G3BP1/G3BP2 double KO (GFP-DDG3BP1/2) | (Kedersha et al., 2016) |
| U2OS-GFP-G3BP1-G3BP1/G3BP2 double KO (GFP-G3BP1-DDG3BP1/2) | (Kedersha et al., 2016) |
| U2OS-UBAP2L KO (DUBAP2L) | (Sanders et al., 2020) |
| U2OS-G3BP1 KO (DG3BP1) | (Kedersha et al., 2016) |
| U2OS-G3BP2 KO (DG3BP2) | (Kedersha et al., 2016) |
| MDA-MB-468-WT | ATCC |
| MDA-MB-468-G3BP1/G3BP2 KO double (DDG3BP1/2) | (Yang et al., 2024) |

**Table S2.** Details of siRNAs used in this study.

| Target | siRNA | Source |
|---|---|---|
| MYH9 (Human) Entrez Gene 4627 | siGENOME Human MYH9 siRNA – SMARTpool, targeting:<br><br>GUAUCAAUGUGACCGAUUU<br><br>CAAAGGAGCCCUGGCGUUA<br><br>GGAGGAACGCCGAGCAGUA<br><br>CGAAGCGGGUGAAAGCAAA | Dharmacon (A Horizon Discovery Group Company) Discovery, cat no. L-007668-00 |
| Non-Targeting control | siGENOME Non-Targeting siRNA Pool #2, targeting:<br><br>UAAGGCUAUGAAGAGAUAC<br><br>AUGUAUUGGCCUGUAUUAG<br><br>AUGAACGUGAAUUGCUCAA<br><br>UGGUUUACAUGUCGACUAA | Horizon Discovery, cat no. D-001206-14 |

**Table S3.** Information for antibodies used in the study.

| Antigen | Host species | Catalog number | Source | IF dilution | WB dilution |
|---|---|---|---|---|---|
| G3BP1 | Mouse | sc-365338 | Santa Cruz Biotechnology, Inc. | 1:200 | 1:1000 |
| eIF3b | Rabbit | PA5-117928 | Invitrogen | 1:200 | N/A |
| MYH9 | Mouse | 60233-1-Ig | Protein tech | N/A | 1:1000 |
| b-actin | Mouse | 66009-1-Ig | Protein tech | N/A | 1:2000 |
| G3BP1 | Rabbit | A302-034A | Bethyl Labs | 1:200 | N/A |
| HEDLS/ P70 S6 kinase | Mouse | sc-8418 | Santa Cruz Biotechnology, Inc. | 1:200 | N/A |
| UBAP2L | Rabbit | A300-534A | Bethyl Labs | N/A | 1:1000 |
| DDX6 | Rabbit | A300-461A | Bethyl Labs | N/A | 1:1000 |
| EDC3 | Mouse | sc-365024 | Santa Cruz Biotechnology, Inc. | N/A | 1:1000 |
| G3BP2 | Rabbit | A302-040A | Bethyl Labs | N/A | 1:1000 |
| USP10 | Mouse | sc-365828 | Santa Cruz Biotechnology, Inc. | N/A | 1:500 |
| Caprin1 | Rabbit | 15112-1-AP | Protein tech | N/A | 1:1000 |
| G3BP1 | Mouse | sc-365338 | Santa Cruz Biotechnology, Inc. | 1:500 | 1:1000 |
| p-eIF2a | | sc-8418 | Santa Cruz Biotechnology, Inc. | 1:200 | N/A |
| Total eIF2a | | 10970 | Protein Tech | 1:200 | N/A |
| puromycin | | sc-1751 | Santa Cruz Biotechnology, Inc. | 1:500 | N/A |
| Cy3 AffiniPure™ Goat Anti-Rabbit IgG (H + L) | | 111-165-003 | Jackson Immuno | 1:2000 | |
| Cy™5 AffiniPure Goat Anti-Mouse IgG (H + L) | | 115-005-003 | Jackson ImmunoResearch | 1:200 | |
| Cy™2 AffiniPure Goat Anti-Rabbit IgG (H + L) | | 111-005-003 | Jackson ImmunoResearch | 1:200 | |
| Alexa Fluor® 488 AffiniPure® Goat Anti-Mouse IgG, $F(ab')_2$ fragment specific | | 115-545-006 | Jackson ImmunoResearch | 1:200 | |
| Peroxidase-Conjugated AffiniPure Donkey anti-Mouse | Donkey | 715-035-150 | Jackson ImmunoResearch | NA | 1:5000 |
| Peroxidase-Conjugated AffiniPure Donkey anti-Goat) at 1:5000 | Donkey | 711-005-152 | Jackson ImmunoResearch | NA | 1:5000 |

### Supplementary Figure 1.

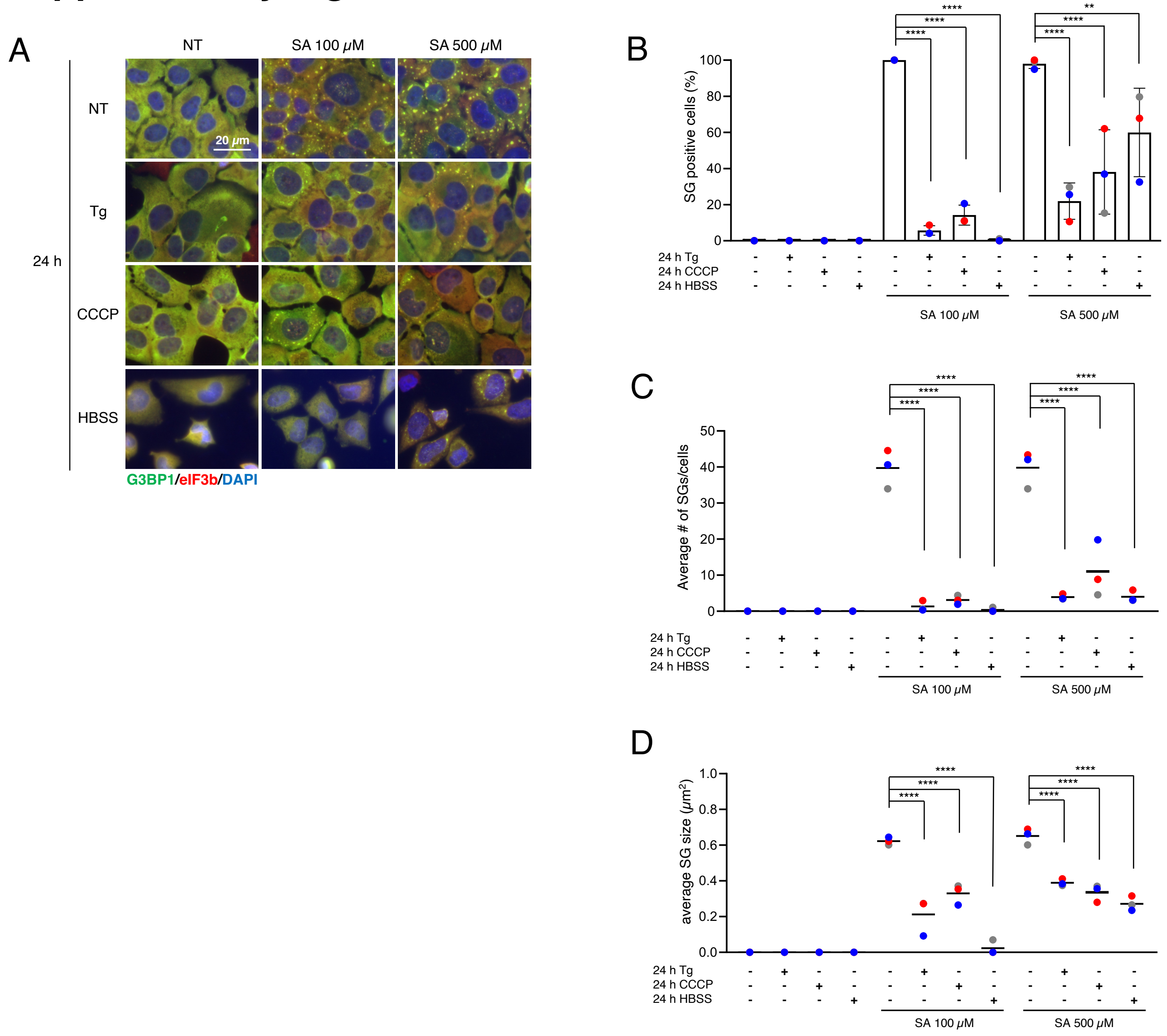


**Supplementary Figure 1. Chronic incubation of Tg, CCCP, and HBSS also decreases the formation and size of SG.** (A–D) U2OS cells were subjected to treatment with 100 or 500 μM SA for 1 h after preincubation with 1 μM Tg, 60 μM CCCP or HBSS for 24 h. Unstressed cells (NT) were used as a control. (A) Representative images of U2OS cells stained with G3BP1 (green), eIF3B (red), and DAPI (blue) after the cells had been subjected to specific stresses. Scale bar = 20 μm. (B) Cells were examined for the presence of the core SG markers G3BP1, and eIF3B. (C) Cells were examined for the average number of G3BP1 per cell. (D) Cells were examined for the average size of G3BP1. (B–D) P values were assessed using a one-way ANOVA (p** < 0.01, p**** < 0.0001) Results are mean ± S.E.M. (n = 12).

### Supplementary Figure 2.

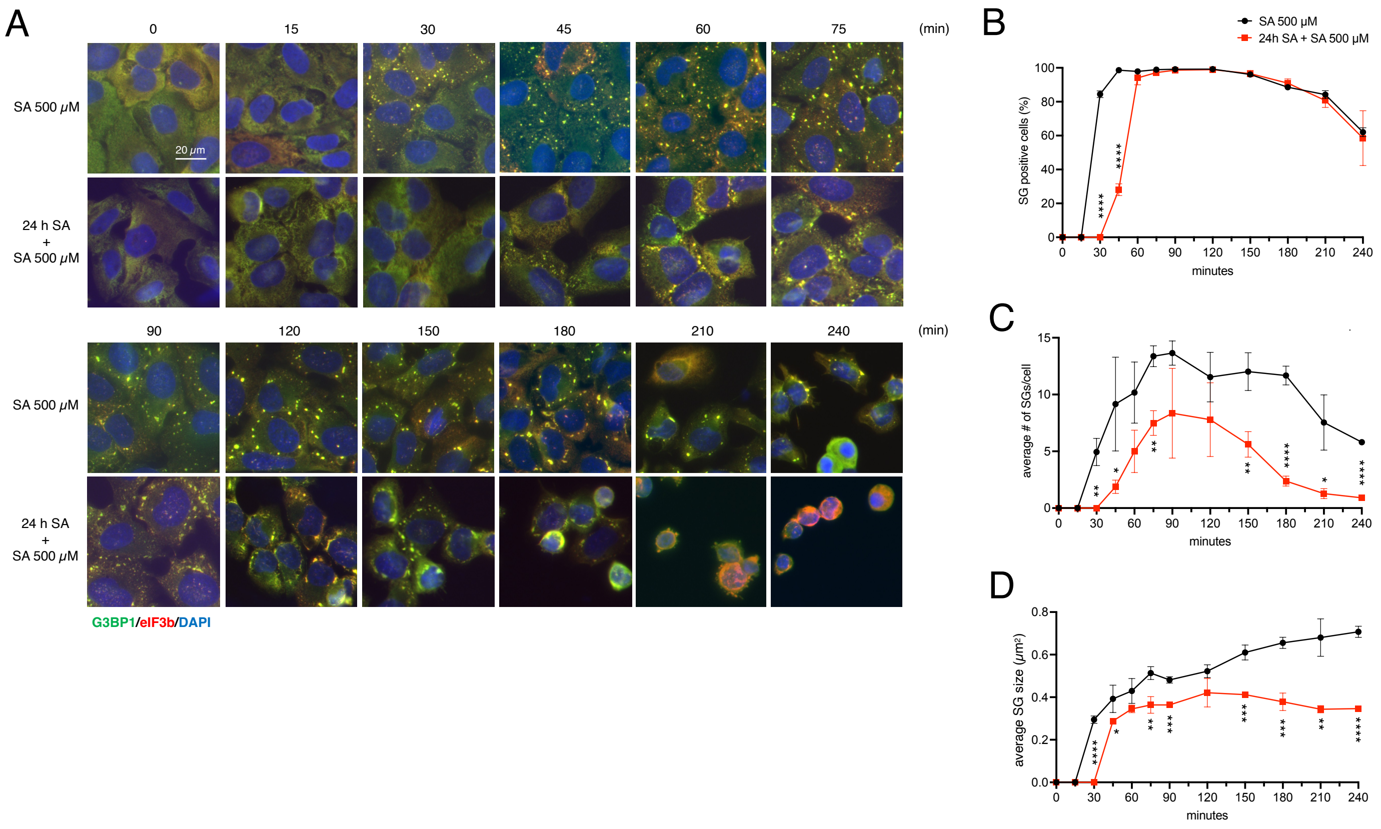


**Supplementary Figure 2. Chronic stress negatively regulates the size of SGs at any time points of SG induction**

(A–D) U2OS cells were subjected to treatment with 500 μM SA for 15, 30, 45, 60, 75, 90, 120, 150, 180, 210, 240 min after preincubation with 10 μM SA for 24 h. Unstressed cells (NT) were used as a control. (A) Representative images of U2OS cells stained with G3BP1 (green), eIF3B (red), and DAPI (blue). Scale bar = 20 μm. (B) Cells were examined for the presence of the core SG markers G3BP1, and eIF3B. (C) Cells were examined for the average number of G3BP1 per cell. (D) Cells were examined for the average size of G3BP1. (B–D) Four images were taken for each of the three biological replicates (independent experiments). P values were assessed using a one-way ANOVA ($p^* < 0.05$, $p^{**} < 0.01$, $p^{***} < 0.001$, $p^{****} < 0.0001$). Results are mean ± S.E.M. (n = 3).

### Supplementary Figure 3.

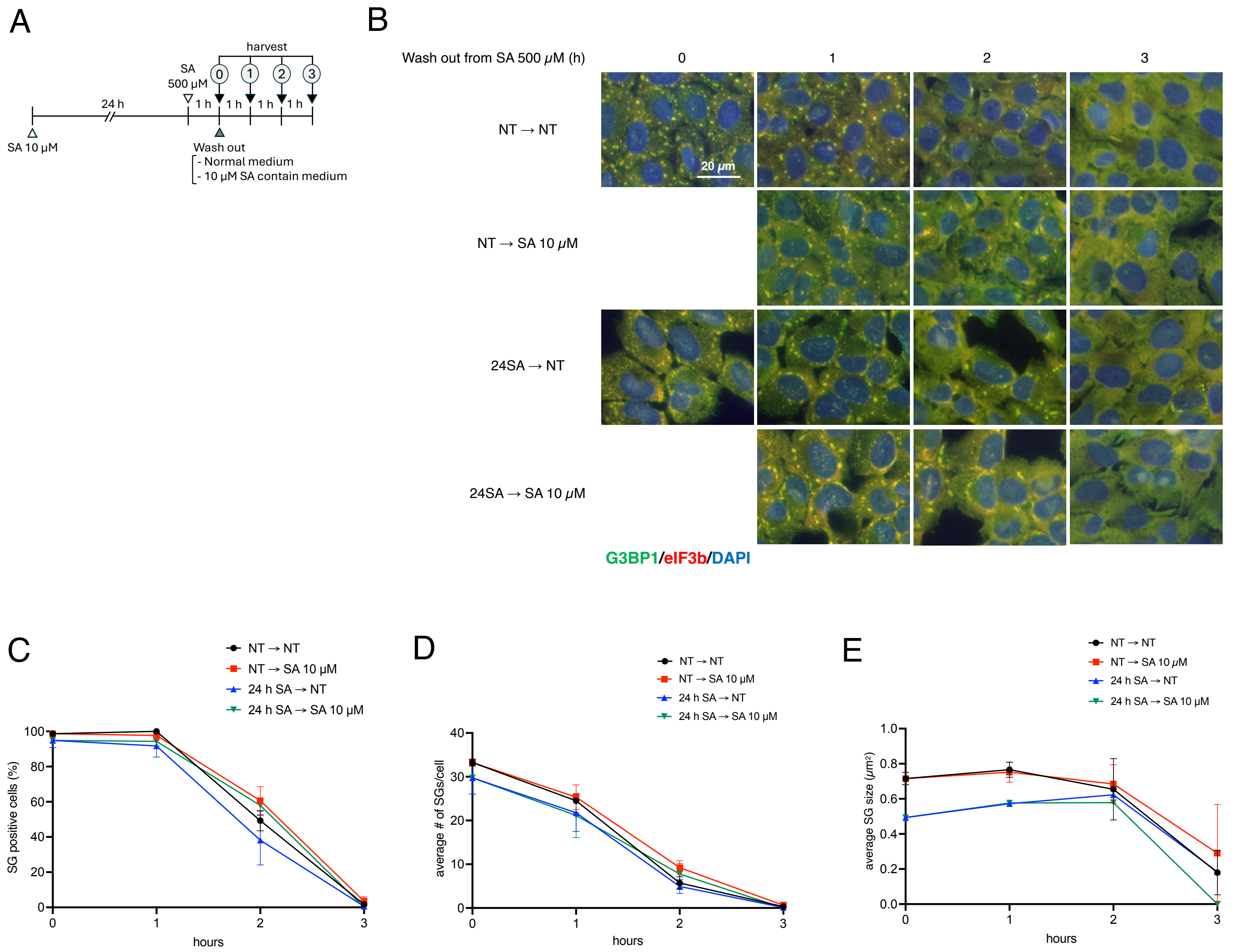


**Supplementary Figure 3. Chronic stress does not affect the disassembly of SG**

U2OS cells were subjected to treatment with 500 μM SA for 1 h after preincubation with 10 μM SA for 24 h prior to changing the medium to fresh or 10 μM SA contained medium. Unstressed cells (NT) were used as a control. (A) Schematic illustration of the experimental timeline. (B) Representative images of U2OS cells stained with G3BP1 (green), eIF3B (red), and DAPI (blue). (C) Cells were examined for the presence of the core SG markers G3BP1, and eIF3B. Scale bar = 20 μm. (D) Cells were examined for the average number of G3BP1 per cell. (E) Cells were examined for the average size of G3BP1. (C–E) Results are mean ± S.E.M. (n = 3).

### Supplementary Figure 4.

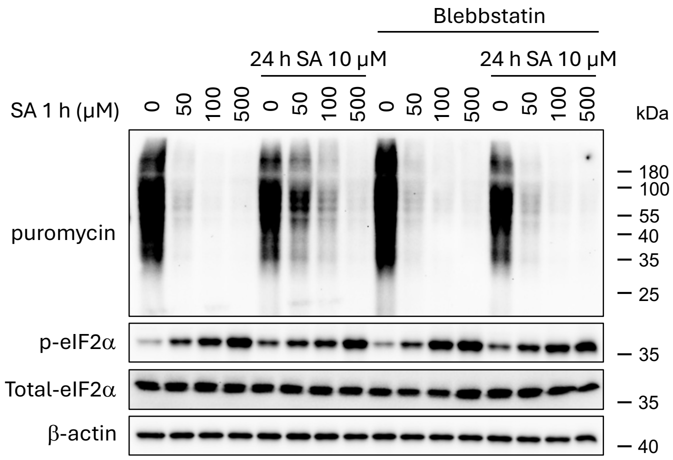


**Supplementary Figure 4. Blebbstatin does not rescue translation under chronic stress**

U2OS cells were subjected to treatment with 50, 100, or 500 μM SA for 1 h after pre-incubation with 10 μM SA and/or 100 μM of blebbstatin for 24 h. Unstressed cells (NT; SA 0 μM) were used as a control. Cells were pulsed with puromycin and emetine for 5 min and lysed. Cell lysates were subjected to western blotting using antibodies for Puromycin, p-eIF2α, total eIF2α, and β-actin.

### Supplementary Figure 5.

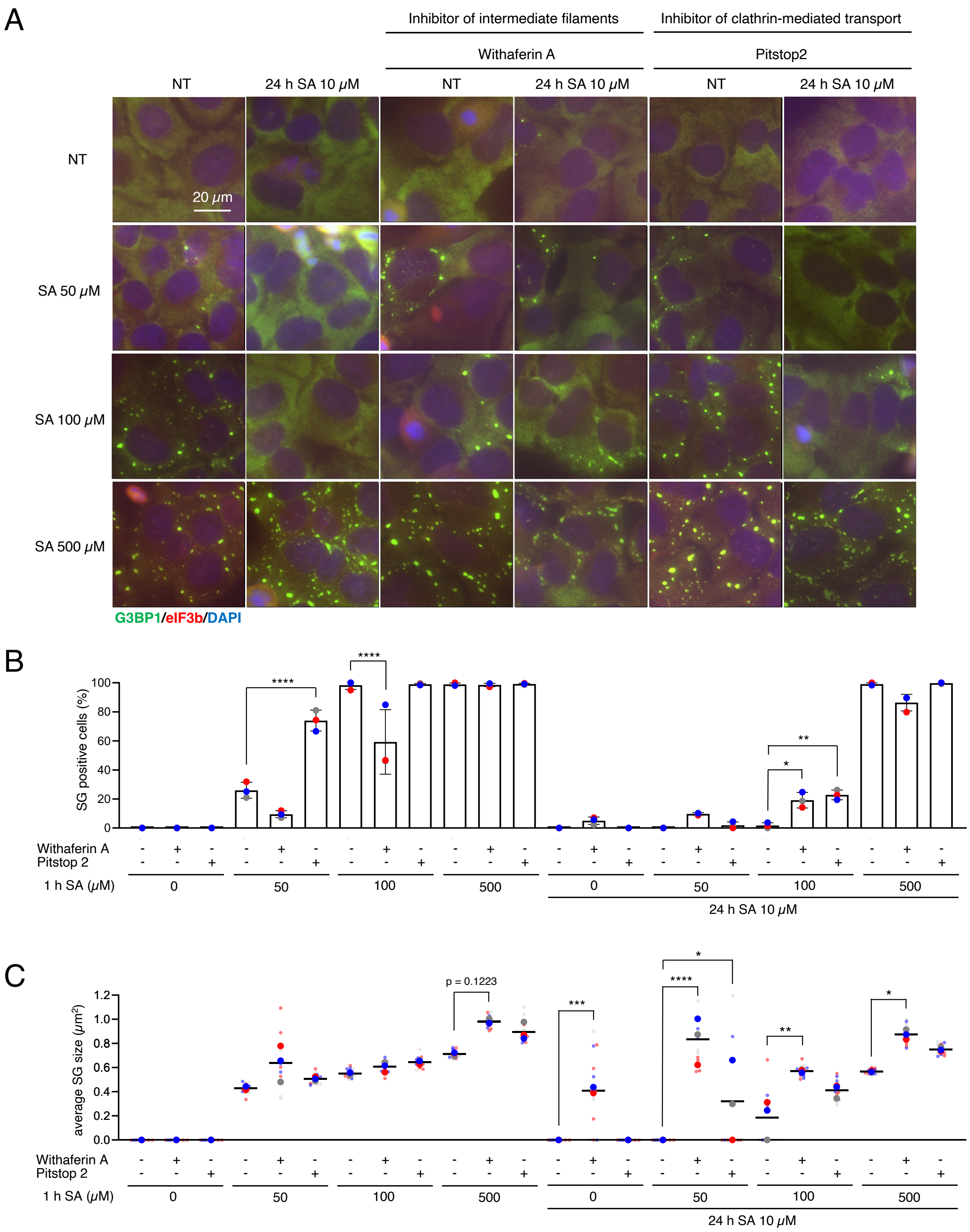


**Supplementary Figure 5. Inhibitors of intermediate filaments and Clathrin-mediated transport positively regulate the SGs**
(A–C) U2OS cells were subjected to treatment with 50, 100, or 500 μM SA for 1 h after preincubation with 10 μM SA and/or 1 μM withaferin A and 30 μM Pitstop 2 for 24 h. Unstressed cells (NT) were used as a control. (A) Representative images of U2OS cells stained with G3BP1 (green), eIF3B (red), and DAPI (blue). Scale bar = 20 μm. (B) Cells were examined for the presence of the core SG markers G3BP1, and eIF3B. (C) Cells were examined for the average size of G3BP1. (B–C) Four images were taken for each of the three biological replicates (independent experiments). Large circle dots represent the mean results from the images (n = 4 technical replicates) of one biological replicate, the same colors of dots are from the same independent experiment. Small circle dots in different shades of the same color are from the same independent experiment. P values were assessed using a one-way ANOVA ($p^* < 0.05$, $p^{**} < 0.01$, $p^{***} < 0.001$, $p^{****} < 0.0001$). Results are mean ± S.E.M. (n = 3).

### Supplementary Figure 6.

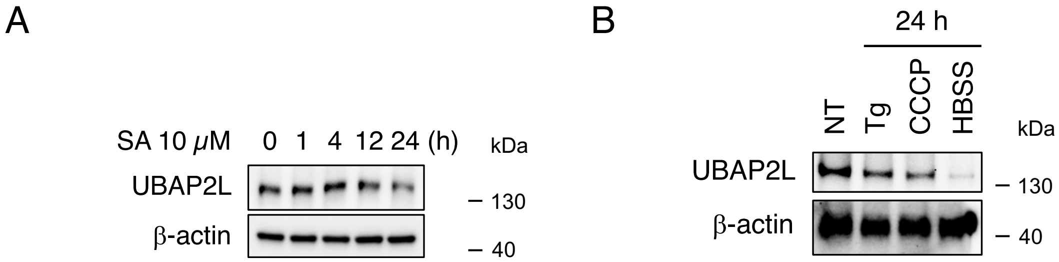


**Supplementary Figure 6. Chronic stresses decrease the protein level of UBAP2L**

(A) U2OS cells were subjected to treatment with 10 μM SA for 0, 1, 4, 12, and 24 h. (B) U2OS cells were subjected to treatment with 1 μM Tg, 60 μM CCCP, or HBSS for 24 h. (A–B) Cell lysates were subjected to western blotting using antibodies for UBAP2L and β-actin.

### Supplementary Figure 7.

A

| | | | | | | |
|---|---|---|---|---|---|---|
| 1 | MAQQAADKYL | YVDKNFINNP | LAQADWAAKK | LVWVPSDKSG | FEPASLKEEV | 50 |
| 51 | GEEAIVELVE | NGKKVKVNKD | DIQKMNPPKF | SKVEDMAELT | CLNEASVLHN | 100 |
| 101 | LKERYYSGLI | YTYSGLFCVV | INPYKNLPIY | SEEIVEMYKG | KKRHEMPPHI | 150 |
| 151 | YAITDTAYRS | MMQDREDQSI | LCTGESGAGK | TENTKKVIQY | LAYVASSHKS | 200 |
| 201 | KKDQGELERQ | LLQANPILEA | FGNAKTVKND | NSSRFGKFIR | INFDVNGYIV | 250 |
| 251 | GANIETYLLE | KSRAIRQAKE | ERTFHIFYYL | LSGAGEHLKT | DLLLEPYNKY | 300 |
| 301 | RFLSNGHVTI | PGQQDKDMFQ | ETMEAMRIMG | IPEEEQMGLL | RVISGVLQLG | 350 |
| 351 | NIVFKKERNT | DQASMPDNTA | AQKVSHLLGI | NVTDFTRGIL | TPRIKVGRDY | 400 |
| 401 | VQKAQTKEQA | DFAIEALAKA | TYERMFRWLV | LRINKALDKT | KRQGASFIGI | 450 |
| 451 | LDIAGFEIFD | LNSFEQLCIN | YTNEKLQQLF | NHTMFILEQE | EYQREGIEWN | 500 |
| 501 | FIDFGLDLQP | CIDLIEKPAG | PPGILALLDE | ECWFPKATDK | SFVEKVMQEQ | 550 |
| 551 | GTHPKFQKPK | QLKDKADFCI | IHYAGKVDYK | ADEWLMKNMD | PLNDNIATLL | 600 |
| 601 | HQSSDKFVSE | LWKDVDRIIG | LDQVAGMSET | ALPGAFKTRK | GMFRTVGQLY | 650 |
| 651 | KEQLAKLMAT | LRNTNPNFVR | CIIPNHEKKA | GKLDPHLVLD | QLRCNGVLEG | 700 |
| 701 | IRICRQGFPN | RVVFQEFRQR | YEILTPNSIP | KGFMDGKQAC | VLMIKALELD | 750 |
| 751 | SNLYRIGQSK | VFFRAGVLAH | LEEERDLKIT | DVIIGFQACC | RGYLARKAFA | 800 |
| 801 | KRQQQLTAMK | VLQRNCAAYL | KLRNWQWWRL | FTKVKPLLQV | SRQEEEMMAK | 850 |
| 851 | EEELVKVREK | QLAAENRLTE | METLQSQLMA | EKLQLQEQLQ | AETELCAEAE | 900 |
| 901 | ELRARLTAKK | QELEEICHDL | EARVEEEEER | CQHLQAEKKK | MQQNIQELEE | 950 |
| 951 | QLEEEESARQ | KLQLEKVTTE | AKLKKLEEEQ | IILEDQNCKL | AKEKKLLEDR | 1000 |
| 1001 | IAEFTTNLTE | EEEKSKSLAK | LKNKHEAMIT | DLEERLRREE | KQRQELEKTR | 1050 |
| 1051 | RKLEGDSTDL | SDQIAELQAQ | IAELKMQLAK | KEEELQAALA | RVEEEAAQKN | 1100 |
| 1101 | MALKKIRELE | SQISELQEDL | ESERASRNKA | EKQKRDLGEE | LEALKTELED | 1150 |
| 1151 | TLDSTAAQQE | LRSKREQEVN | ILKKTLEEEA | KTHEAQIQEM | RQKHSQAVEE | 1200 |
| 1201 | LAEQLEQTKR | VKANLEKAKQ | TLENERGELA | NEVKVLLQGK | GDSEHKRKKV | 1250 |
| 1251 | EAQLQELQVK | FNEGERVRTE | LADKVTKLQV | ELDNVTGLLS | QSDSKSSKLT | 1300 |
| 1301 | KDFSALESQL | QDTQELLQEE | NRQKLSLSTK | LKQVEDEKNS | FREQLEEEEE | 1350 |
| 1351 | AKHNLEKQIA | TLHAQVADMK | KKMEDSVGCL | ETAEEVKRKL | QKDLEGLSQR | 1400 |
| 1401 | HEEKVAAYDK | LEKTKTRLQQ | ELDDLLVDLD | HQRQSACNLE | KKQKKFDQLL | 1450 |
| 1451 | AEEKTISAKY | AEERDRAEAE | AREKETKALS | LARALEEAME | QKAELERLNK | 1500 |
| 1501 | QFRTEMEDLM | SSKDDVGKSV | HELEKSKRAL | EQQVEEMKTQ | LEELEDELQA | 1550 |
| 1551 | TEDAKLRLEV | NLQAMKAQFE | RDLQGRDEQS | EEKKKQLVRQ | VREMEAELED | 1600 |
| 1601 | ERKQRSMAVA | ARKKLEMDLK | DLEAHIDSAN | KNRDEAIKQL | RKLQAQMKDC | 1650 |
| 1651 | MRELDDTRAS | REEILAQAKE | NEKKLKSMEA | EMIQLQEELA | AAERAKRQAQ | 1700 |
| 1701 | QERDELADEI | ANSSGKGALA | LEEKRRLEAR | IAQLEEELEE | EQGNTELIND | 1750 |
| 1751 | RLKKANLQID | QINTDLNLER | SHAQKNENAR | QQLERQNKEL | KVKLQEMEGT | 1800 |
| 1801 | VKSKYKASIT | ALEAKIAQLE | EQLDNETKER | QAACKQVRRT | EKKLKDVLLQ | 1850 |
| 1851 | VDDERRNAEQ | YKDQADKAST | RLKQLKRQLE | EAEEEAQRAN | ASRRKLQREL | 1900 |
| 1901 | EDATETADAM | NREVSSLKNK | LRRGDLPFVV | PRRMARKGAG | DGSDEEVDGK | 1950 |
| 1951 | ADGAEAKPAE | | | | | 2000 |

B

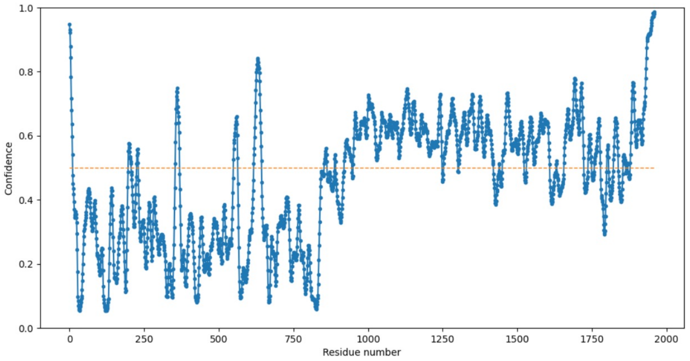


**Supplementary Figure 7. Predicted disordered region of MYH9 protein**

Disordered region of human MYH9 protein amino acids was predicted by protein disorder prediction system (PrDOS).
(A) All amino acids of the human MYH9 protein. Predicted disordered amino acids were red colored. (B) Schematic plots of the human MYH9 amino acids with the confidence of predicted disordered amino acids.
